# Transdifferentiation of epithelial cells and fibroblasts induced by IL-1β fuels neutrophil recruitment in chronic rhinosinusitis

**DOI:** 10.1101/2024.01.28.576762

**Authors:** Xinyu Xie, Pin Wang, Min Jin, Yue Wang, Lijie Qi, Changhua Wu, Shu Guo, Changqing Li, Xiaojun Zhang, Ye Yuan, Xinyi Ma, Fangying Liu, Weiyuan Liu, Heng Liu, Chen Duan, Ping Ye, Xuezhong Li, Larry Borish, Wei Zhao, Xin Feng

**Affiliations:** Department of Otorhinolaryngology, Qilu Hospital of Shandong University, National Health Commission Key Laboratory of Otorhinolaryngology (Shandong University), Jinan, China; Department of Anesthesiology, Qilu Hospital of Shandong University, Jinan, China; Department of Gastroenterology, Qilu Hospital of Shandong University, Jinan, China; Departments of Medicine and Microbiology, University of Virginia Health System, Charlottesville, Virginia, USA; Key Laboratory for Experimental Teratology of the Chinese Ministry of Education, and Key Laboratory of Infection and Immunity of Shandong Province, School of Basic Medical Science, Qilu Hospital, Cheeloo College of Medicine, Shandong University, Jinan, Shandong, China

## Abstract

Neutrophilic inflammation contributes to multiple chronic inflammatory airway diseases, including asthma and chronic rhinosinusitis with nasal polyps (CRSwNP), and is associated with an unfavorable prognosis. Here, using single-cell RNA sequencing (scRNA-seq) to profile human nasal mucosa obtained from the inferior turbinates, middle turbinates, and nasal polyps of CRSwNP patients, we identified two IL-1 signaling-induced cell subsets—*LY6D*^+^ club cells and *IDO1*^+^ fibroblasts—that promote neutrophil recruitment by respectively releasing S100A8/A9 and CXCL1/2/3/5/6/8 into inflammatory regions. IL-1β, a pro-inflammatory cytokine involved in IL-1 signaling, induces the transdifferentiation of *LY6D*^+^ club cells and *IDO1*^+^ fibroblasts from primary epithelial cells and fibroblasts, respectively. In an LPS-induced neutrophilic CRSwNP mouse model, blocking IL-1β activity with a receptor antagonist significantly reduced the numbers of *LY6D*^+^ club cells and *IDO1*^+^ fibroblasts and mitigated nasal inflammation. This study reveals the roles of two cell subsets in neutrophil recruitment and demonstrates an IL-1-based intervention for mitigating neutrophilic inflammation in CRSwNP.

## Introduction

Neutrophilic inflammation is prevalent in multiple chronic inflammatory airway diseases such as asthma, chronic obstructive pulmonary disease, and chronic rhinosinusitis (CRS), and elevated neutrophilic inflammation is positively correlated with adverse patient outcomes^1,2^. CRS is a chronic disorder characterized by inflammation of the nasal mucosa and paranasal sinuses that affects 5-12% of the global adult population^3^. Patients with CRS and nasal polyps (CRSwNP) experience more severe clinical symptoms than those without nasal polyps^4^. Although CRSwNP exhibits a significant association with type 2 inflammation which is characterized by an immune response involving eosinophils^5^, the presence of a neutrophilic inflammation in CRSwNP has been demonstrated in a growing number of patients, and is considered to be associated with glucocorticosteroid resistance, a higher risk of recurrence after surgery, and worse disease outcomes^6^. However, neutrophilic inflammation has been relatively little studied and therapeutic strategies targeting neutrophilic inflammation are currently insufficient in CRSwNP.

Multiple factors drive the neutrophilic inflammation in CRSwNP. CXC chemokines including CXCL1, CXCL2, and CXCL8 are chemotactic factors that guide the neutrophils to the site of inflammation^7^. In a multi-center study, the concentrations of CXCL8 was shown to be greater in NP tissues than that in control tissues, indicating its role in neutrophil recruitment of CRSwNP^8^. Increased protein levels of S100A8, S100A9, and S100A8/A9, were demonstrated in the nasal polyp tissues of CRSwNP patients compared to those in the IT tissues of controls, suggesting evident neutrophil recruitment in CRSwNP^9^. Previous studies have demonstrated that cytokines derived from epithelial cells and stromal cells facilitate neutrophilic inflammation in CRS^10,11^. Nevertheless, specific cell types that secrete these factors and drive neutrophilic inflammation in CRSwNP remain ill-defined.

Here, seeking to identify epithelial and stromal cell subsets that contribute to neutrophilic inflammation in CRSwNP, we profiled human nasal mucosa obtained from the middle turbinates (MTs), inferior turbinates (ITs), and nasal polyps (NPs) of CRSwNP patients and healthy individuals using single-cell RNA sequencing (scRNA-seq). After identifying contributions from *LY6D*^+^ club cells and *IDO1*^+^ fibroblasts, we demonstrated their ability to facilitate neutrophil recruitment in cells stimulated with IL-1β, including primary fibroblasts and air-liquid interface (ALI) cultures developed from primary nasal epithelial cells. Blocking the activity of IL-1β attenuated nasal inflammation in an LPS-induced neutrophilic CRSwNP mouse model. These findings uncover the cell types that drive neutrophilic inflammation in CRSwNP, and highlight potential therapeutic agents targeting IL-1β as interventions against neutrophilic CRSwNP.

## Results

### Single-cell profiling of nasal mucosa from multiple anatomical regions in CRSwNP patients identifies diverse disease-specific cell subsets

We initially profiled the CRSwNP cell type landscape by preparing freshly dissociated samples of middle turbinate (MT), inferior turbinate (IT), and nasal polyp (NP) tissues from CRSwNP patients and healthy individuals and obtaining full-length scRNA-seq profiles (Fig. 1a). Inferior turbinates have been used as control tissues for nasal polyps in previous studies^12,13^. Most NPs originate from the ethmoid sinuses located around the MT tissues, and MT tissue removal has been shown to reduce the recurrence of NPs in refractory CRS^14^. We therefore selected MT, IT, and NP tissues to compare differences in their cellular composition in an inflammatory milieu. Unsupervised clustering divided the 219,716 cells that passed strict quality-control into six compartments with conserved signatures, including epithelial cells, T/innate lymphoid cells (ILCs), B/plasma cells, mononuclear phagocytes/dendritic cells (MNPs/DCs), mast cells, and stromal cells (Fig.1b-d, Extended Data Fig. 1a-d and Extended Data Fig. 2). B/plasma cells, MNPs/DCs, and mast cells, were barely detectable in the IT tissues from healthy individuals, supporting an extensive inflammatory milieu in both nasal polyps and nasal mucosa of CRSwNP patients, regardless of the anatomical regions in which they occur (Fig. 1b, e). Of note, each of the subsets contained cells from each sample, indicating that the cell lineages and expression status were consistent throughout samples and did not represent sample-specific subpopulations or batch effects (Extended Data Fig. 3a, b).

**Fig. 1:**
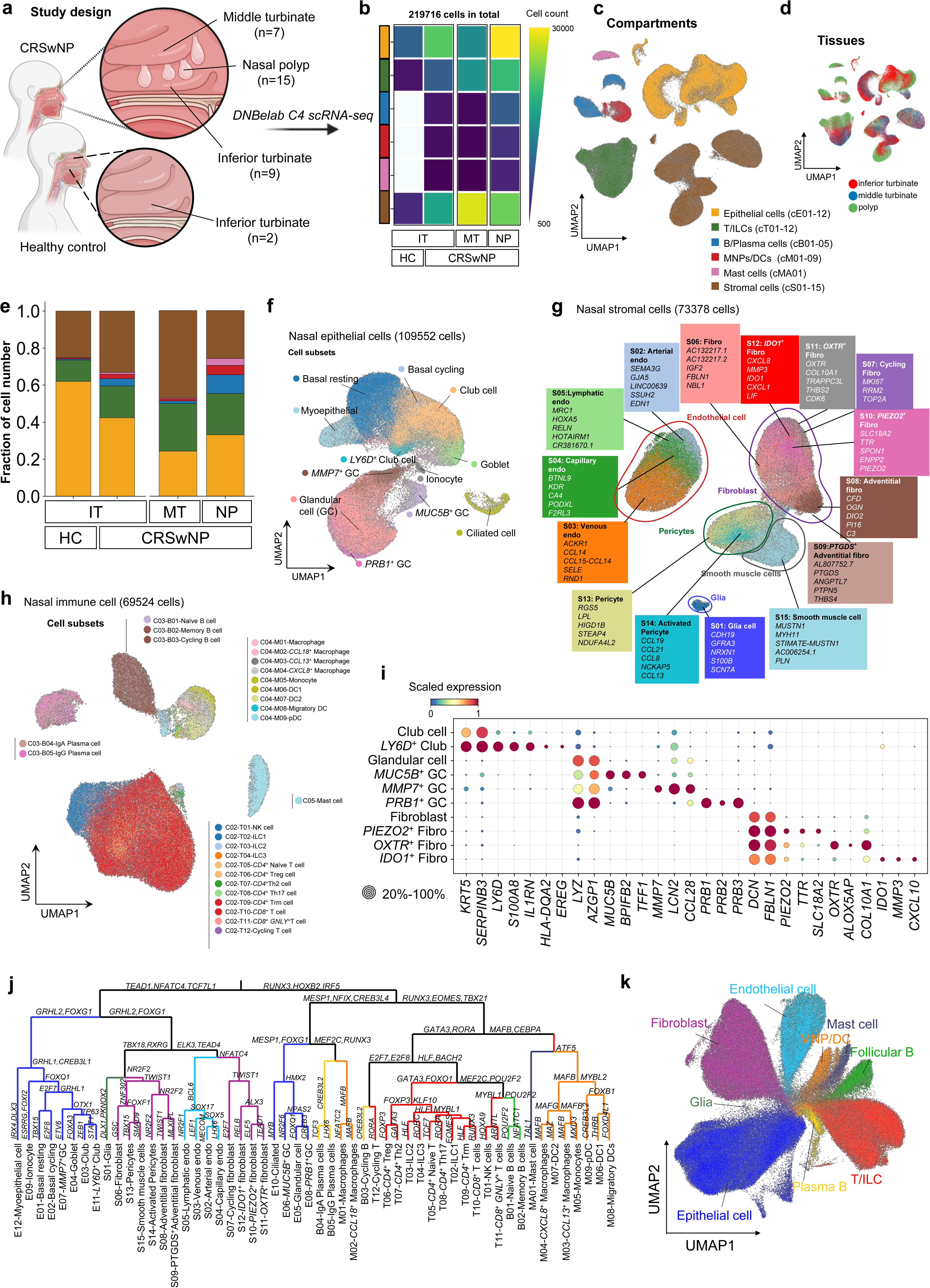
Analysis of middle turbinates, inferior turbinates, and nasal polyps from CRSwNP patients and healthy individuals. **a** For the study design, 33 samples were collected from distinct anatomical regions (inferior turbinates, middle turbinates, and nasal polyps) of CRSwNP patients and healthy individuals. scRNA-seq (DNBelab C4) acquired 219,716 high-quality cells. Created with BioRender.com. **b** Cell counts by anatomical region for each compartment. The colors of the cell compartments are consistent in panel (**b**) and (**c**). **c** Unsupervised sub-clustering preliminarily divided the cells into six compartments. **d** UMAP (uniform manifold approximation and projection) embedding by three anatomical regions. **e** Bar plot depicting the cell compositions of the indicated anatomical regions of human nasal mucosa from CRSwNP patients and healthy individuals. The colors of the cell compartments are consistent in panel (**b**) and (**c**). **f** UMAP displaying typical cell subsets of the nasal mucosal epithelium. **g** UMAP displaying 15 cell subsets of stromal cells in the nasal mucosa with the gene signatures of each subset indicated in the colored boxes. **h** UMAP displaying immune cell subsets in all samples. **i** Bubble heatmap showing marker genes across cell subsets of interest in this study. **j** A dendrogram of regulons for all cell subsets constructed from the fate decision tree analysis. TFs at each branching point are representative regulons of subjacent groups. The colors of the cell subsets are consistent in panel (**j**) and (**k**). **k** UMAP showing six cell compartments and some cell subsets based on the regulons from the fate decision tree analysis presented in panel (**j**). The colors of the cell subsets are consistent in panel (**j**) and (**k**).

To identify cell subsets associated with inflammation regulation, we performed unsupervised clustering based on marker genes on the epithelial cell compartment, which revealed seven cell types annotated as basal cells, myoepithelial cells, club cells, goblet cells, ciliated cells, ionocytes, and glandular cells (GCs) (Fig. 1f). Among the identified subsets, *LY6D*^+^ club cells have not been reported, while the *PRB1^+^* GC and *MUC5B*^+^ GC subsets were previously observed in the nasal mucosa of CRSwNP patients^15^. Pathway enrichment analysis revealed that *PRB1*^+^ GCs are associated with erythrocyte renewal and metabolism, while *MUC5B*^+^ GCs are involved in protein glycosylation, especially mucin glycosylation (Extended Data Fig. 4a-d). The stromal cell compartment was divided into five cell types (endothelial cells, pericytes, fibroblasts, smooth muscle cells, and glia) and then further classified into 15 yet-finer subsets based on marker gene expression; among these subsets, *PIEZO2*^+^, *IDO1*^+^, and *OXTR*^+^ fibroblasts have not been reported in previous studies of nasal mucosa from CRSwNP patients (Fig. 1g, 1i). Given that *OXTR*^+^ fibroblasts were detected only in NP tissues, these cells may be involved in NPs development (Fig. 4d). The immune cell compartment was subclustered into five cell types, including mast cells, mononuclear phagocytes/dendritic cells (MNPs/DCs), plasma cells, B cells, and T/innate lymphoid cells (T/ILCs), which were subsequently grouped into 27 yet-finer subsets (Fig. 1h and Extended Data Fig. 5a-d). ILC1/2/3 were enriched in NP tissues, reflecting a mixed pattern of inflammation in CRSwNP^16–18^ (Extended Data Fig. 5c).

To demonstrate the relationship between cell subsets during differentiation, we constructed a transcription factor fate decision tree for cells spanning different anatomical regions (Fig. 1j, k). This analysis suggested that transcription factors, such as *STAT1*, *ELF5*, *TEAD1*, and *CREB3*, are regulons modulating the differentiation of different cell subsets, further demonstrating the correctness of the sub-clustering across the samples. Collectively, these findings reveal the cellular heterogeneity in the inflammatory environment across three anatomical regions, and identify disease-specific cell subsets that may regulate immune response in CRSwNP.

### *IDO1*^+^ fibroblasts and *LY6D*^+^ club cells contribute to neutrophil recruitment in CRSwNP

CRSwNP patients exhibit both eosinophilic and neutrophilic inflammation^19^. Increased neutrophilia was detected in the mucosa of NP tissues from CRSwNP patients (Fig. 2a). We also used the xCell algorithm to quantify neutrophil infiltration in a bulk RNA-seq dataset of CRSwNP (GSE179265), and again detected the significantly elevated neutrophilia in CRSwNP samples as compared to healthy tissue samples (Fig. 2b). Seeking to identify epithelial and stromal cell subsets contributing to neutrophil infiltration, we generated an integrated dataset built from our data and another CRSwNP scRNA-seq dataset (HRA000772) (Extended Data Fig. 6a)^20^, and subsequently used an algorithm combining Networkx, Community, and Pygraphviz to plot chemokine-chemokine receptor interaction networks and infer the strongly interacting cell subset pairs. Notable signals from the network included a superlatively strong interaction between *IDO1*^+^ fibroblasts and neutrophils (Fig. 2c), with *MMP7*^+^ GCs and *LY6D*^+^ club cells also interacting strongly with neutrophils (Fig. 2c).

**Fig. 2:**
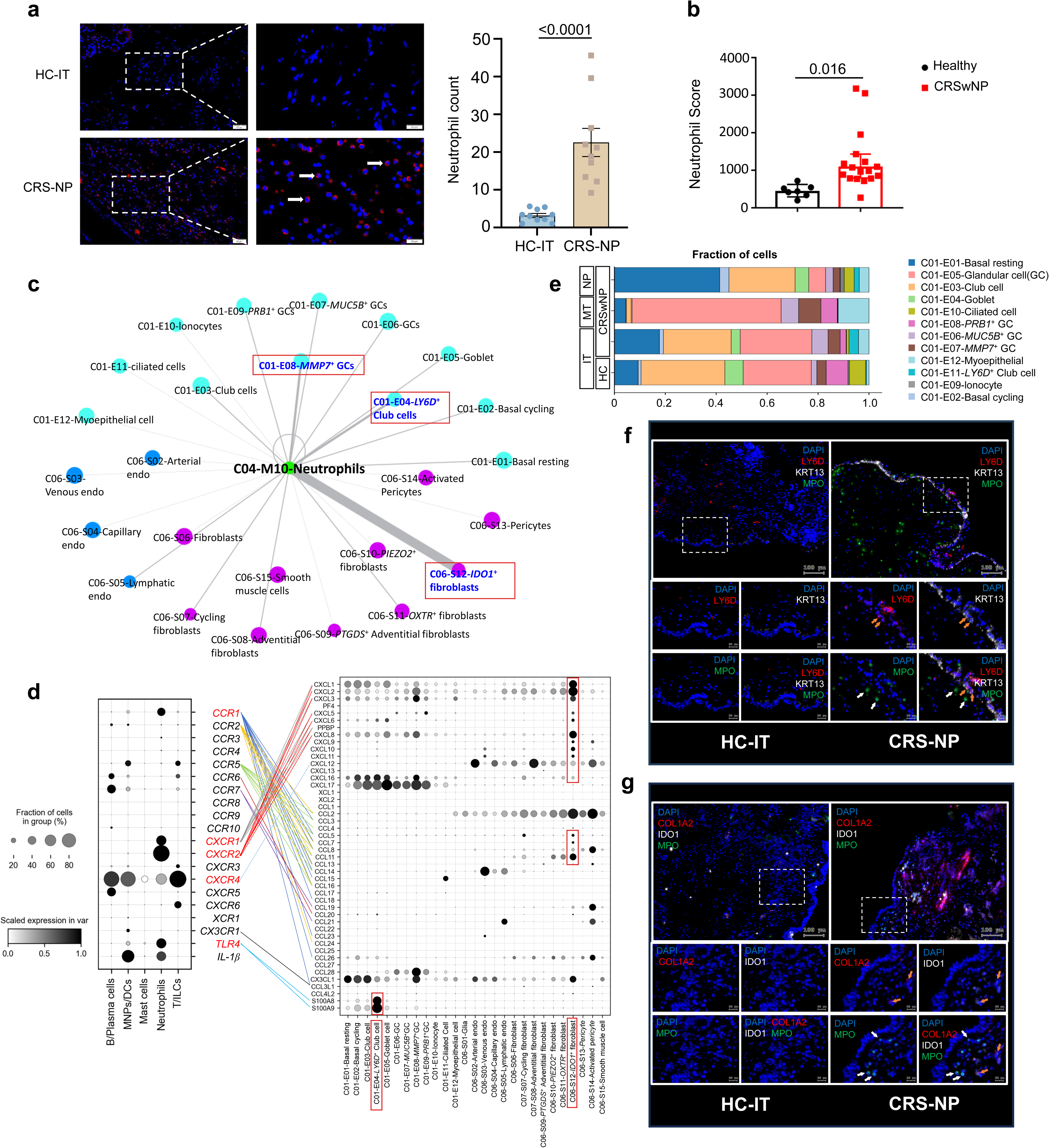
Identification of cell subsets involved in neutrophil recruitment in CRSwNP. **a** Representative image and quantification of immunofluorescence staining for MPO (red) and the nuclear marker DAPI (blue) in IT tissue from healthy individuals (HC-IT) and NP tissue from CRSwNP patients (CRS-NP). Neutrophils are indicated by white arrows. Scale bar, 50 μm (left), 20 μm (right). The data are presented as the means ± SEM. The *P-value* was calculated and reported using the two-tailed Student’s t-test. **b** Neutrophil scores obtained using the xCell algorithm for CRSwNP patients and healthy individuals. The *P-value* was calculated and reported using a two-tailed Student’s t-test. **c** Cell-cell interaction and network representation analysis based on chemokine-chemokine receptor interactions. The nodes with a degree of zero and a connection strength less than the average of all the edges were eliminated. The sizes of the nodes were defined as the log2 (counts+1) of the cell subsets. The thickness of the link reflects the degree of the interaction. Cell subsets that strongly interact with neutrophils are indicated in red boxes. **d** Bubble heatmap for chemokine-chemokine receptor interactions between immune cells and epithelial/stromal cells. Previously validated interactions are indicated by colored straight lines. Chemokines predominantly expressed in *LY6D*^+^ club cells and *IDO1*^+^ fibroblasts are indicated in red boxes. **e** Bar plot depicting the cell composition of epithelial cell subsets in the indicated anatomical regions of human nasal mucosa from CRSwNP patients and healthy individuals. **f** Immunofluorescence staining for LY6D (red), KRT13 (white), MPO (green), and the nuclear marker DAPI (blue) in CRS-NP tissue and HC-IT tissue. The epithelium is indicated with orange arrows. Neutrophils are indicated with white arrows. Scale bar, 20 μm. **g** Immunofluorescence staining for COL1A2 (red), IDO1 (white), MPO (green), and the nuclear marker DAPI (blue) in HC-IT tissue and CRS-NP tissue. The epithelium is indicated with orange arrows. Neutrophils are indicated with white arrows. Scale bar, 20 μm.

In particular, the chemokine receptors enriched in neutrophils (*CCR1* and *CXCR1/2/4*) matched extensively with chemokines highly expressed in *IDO1*^+^ fibroblasts (such as *CXCL1/2/3/5/6/8* and *CCL5/7/8/11*)^21,22^(Fig. 2d). The interaction between neutrophils and *MMP7*^+^ GCs was characterized by high *CXCR2* expression in neutrophils and strong *CXCL2/3* expression in MMP7^+^ GCs. *LY6D*^+^ club cells interacted with neutrophils by expressing high levels of S100A8/A9, and their receptor TLR4 was expressed mainly on neutrophils (Fig. 2d). However, *MMP7*^+^ GCs did not show much difference in proportion of total epithelial cells in different anatomical regions (Fig. 2e). They were probably a subset of cells with an intermediate state based on their low pseudotime ct values calculated by RNA velocity (Extended Data Fig. 4a-d). Therefore, *MMP7*^+^ GCs were not considered to be associated with neutrophil infiltration in CRSwNP.

In contrast to that of *MMP7*^+^ GCs, the proportion of *LY6D*^+^ club cells was greater in the IT and NP tissues of CRSwNP patients than in the IT tissue of healthy individuals (HC-IT), suggesting their potential role in CRSwNP development (Fig. 2e). Consistent with these findings, using the HRA000772 dataset, we also detected higher proportions of *LY6D*^+^ club cells and *IDO1*^+^ cells in the CRSwNP with higher neutrophil numbers (neCRSwNP) than those with lower neutrophil numbers (eCRSwNP) (Extended Data Fig. 6b-d). We then conducted immunofluorescence analyses on NP tissues from 6 CRSwNP patients (CRS-NP) and IT tissues from 6 healthy controls (HC-IT), and the results revealed a preferential distribution of neutrophils (MPO^+^ cells) in the LY6D^+^ club cell-rich and IDO1^+^ fibroblast-rich regions (Fig. 2f, g), supporting their ability to recruit neutrophils in an inflammatory milieu in CRSwNP. These results collectively support that *LY6D*^+^ club cells and *IDO1*^+^ fibroblasts facilitate neutrophil recruitment in CRSwNP.

### *LY6D*^+^ club cells drive IL-1 signaling-mediated neutrophilic inflammation in CRSwNP

We further compared the proportions of *LY6D*^+^ club cells across anatomical regions. An elevated proportion of *LY6D*^+^ club cells within the total epithelial cell population was noted in CRS-ITs compared to HC-ITs, and in NP tissues compared to adjacent MT tissues (Fig. 3a). We then performed immunofluorescence analyses to evaluate the distribution of *LY6D*^+^ club cells in different tissues. Immunostaining detected only a few *LY6D*^+^ cells in normal IT tissues, but more *LY6D*^+^ cells in NP tissues, reflecting the preferential induction of *LY6D*^+^ club cells in an inflammatory milieu (Fig. 3b, c). *LY6D*^+^ club cells were highly conserved across three anatomical regions as indicating in fate decision tree, and *PITX1* ranked as the top differentially expressed transcription factor determining *LY6D*^+^ club cell differentiation (Fig. 3d, e).

**Fig. 3:**
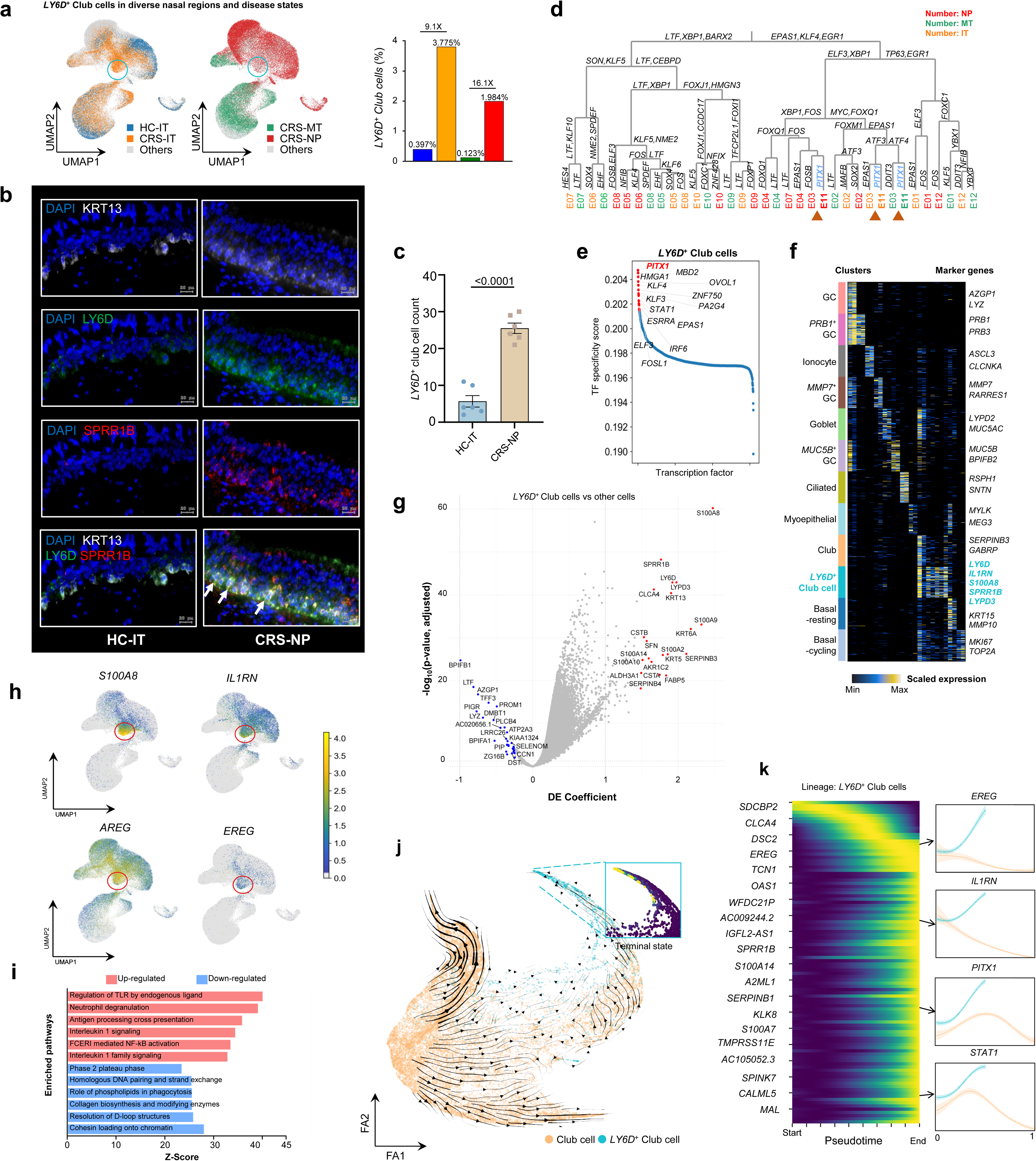
*LY6D*^+^ club cells regulate neutrophilic inflammation in CRSwNP. **a** UMAP displaying the distribution of *LY6D*^+^ club cells in the indicated anatomical regions of human nasal mucosa from CRSwNP patients and healthy individuals. Enriched *LY6D*^+^ club cells are indicated with blue circles (left panel). The proportions of *LY6D*^+^ club cells in the indicated anatomical regions are shown on the right. The colors are consistent in the two panels. **b** Representative immunofluorescence staining for LY6D (green), SPRR1B (red), KRT13 (white, marker gene of club cells) and the nuclear marker DAPI (blue) in HC-IT tissue and CRS-NP tissue. The white arrows indicate colocalization of LY6D, SPRR1B and KRT13. Scale bar, 20 μm. **c** Quantification of the data in panel (**b**). The data are presented as the means ± SEM. The *P-value* was calculated and reported using a two-tailed Student’s t-test. **d** A dendrogram of regulons for epithelial cell subsets in the indicated anatomical regions constructed from the fate decision tree analysis. The transcription factors at each branching point are representative regulons of subjacent groups. The brown triangles show the proximity between *LY6D*^+^ club cells (E11) and club cells (E03) during differentiation. The numbering of cell subsets is consistent with that in Fig. 1(k). **e** TFs enriched in *LY6D*^+^ club cells aligned by TF specificity score. *PITX1* (red) is the top transcription factor responsible for *LY6D*^+^ club cell differentiation. **f** Heatmap of gene expression analyzed by scRNA-seq displaying representative genes for 12 epithelial cell subsets. *LY6D*^+^ club cells are highlighted in cyan letters. **g** Volcano plot displaying the differentially expressed genes (DEGs) between *LY6D*^+^ club cells and other epithelial cell subsets. The *P-value* was calculated and reported using a two-tailed Student’s t-test. **h** UMAP with the epithelial cell compartment displaying the expression of four upregulated genes (IL1RN, S100A8, AREG, and EREG) in *LY6D*^+^ club cells. *LY6D*^+^ club cells are indicated with red circles. **i** Pathway enrichment analysis revealing the enriched signaling pathways in *LY6D*^+^ club cells when compared with those in other epithelial cells. **j** RNA velocity analysis based on RNA splicing information indicating that *LY6D*^+^ club cells are maturely differentiated club cells. **k** Heatmap displaying dynamic changes in the expression of functional genes and TFs during the maturation process of *LY6D*^+^ club cells.

Previous studies have showed that the expression of *S100A8* and *S100A9* is elevated in nasal polyps as compared to control tissues^23,24^, and is associated with neutrophilic inflammation and CRS severity^25^. By exhibiting epithelial cell subset marker gene expression via heatmap, we observed the upregulation of *S100A8* in *LY6D*^+^ club cells (Fig. 3f). We next explored the differentially expressed genes (DEGs) in *LY6D*^+^ club cells as compared to other epithelial cells (Fig. 3g). In addition to *LY6D* and *S100A8*, *S100A9* was also significantly upregulated in *LY6D*^+^ club cells (Fig. 3g). UMAP showed that *LY6D*^+^ club cells were the main cell source of *S100A8* and *S100A9* in the epithelium that may promote neutrophil chemotaxis in CRSwNP^26^ (Fig. 3h and Fig. 5b). The high expression of *EREG* and *AREG* in *LY6D*^+^ club cells indicated their involvement in eosinophil reprogramming and goblet metaplasia in response to inflammation^27,28^ (Fig. 3h).

**Fig. 4:**
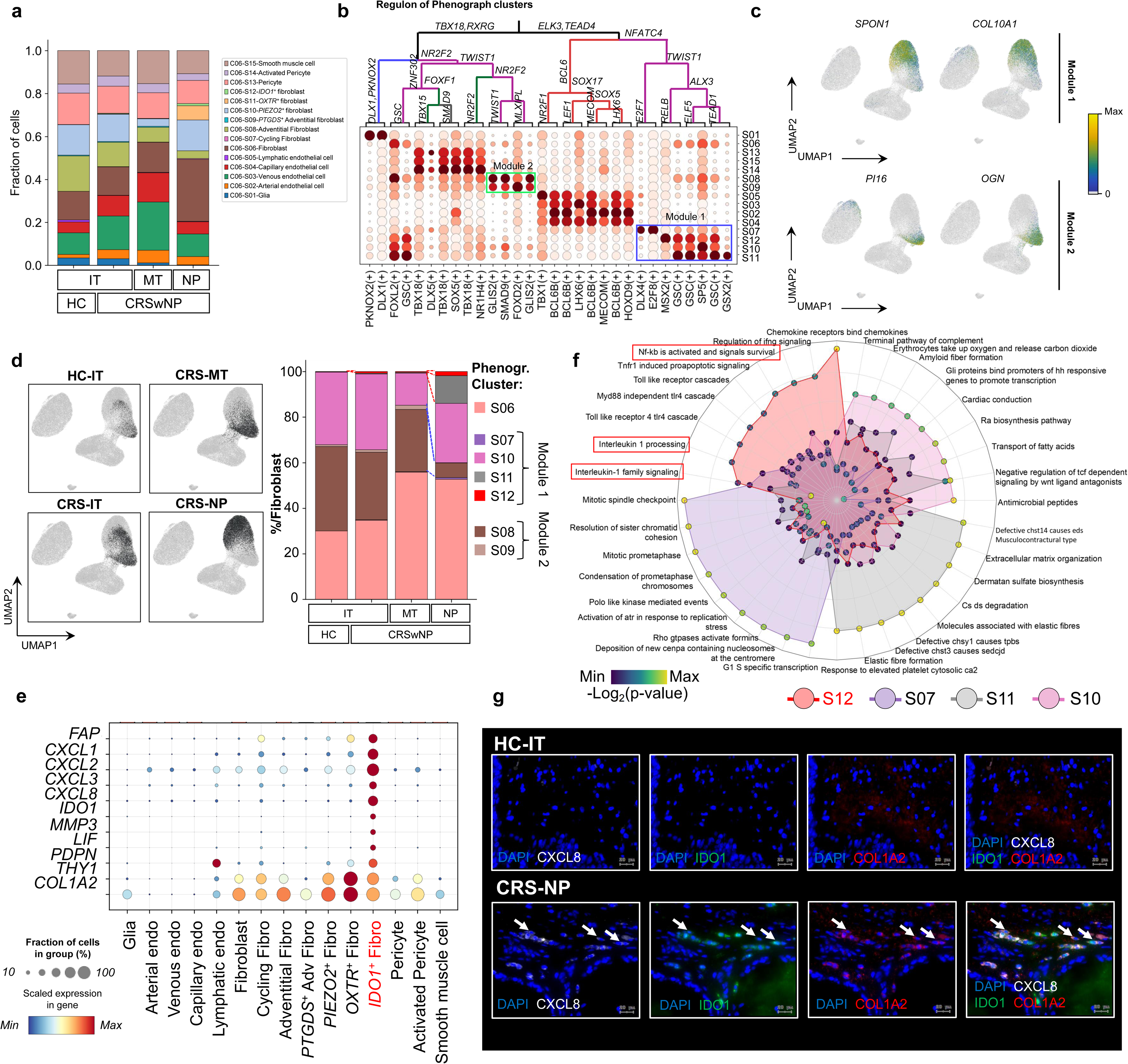
*IDO1*^+^ fibroblasts contribute to IL-1 signaling-mediated neutrophilic inflammation in CRSwNP. **a** Bar plot depicting the cell composition of stromal cell subsets in the indicated anatomical regions of human nasal mucosa from CRSwNP patients and healthy individuals. **b** Transcription factor fate decision tree analysis of stromal cells displaying two distinguishable modules consisting of six fibroblast subsets with remarkable differences in TF patterns. Module 1 and module 2 are indicated by blue and green boxes, respectively. **c** UMAP displaying the expression of marker genes of the two modules in the stromal cell compartment. **d** UMAP of stromal cell compartment displaying fibroblasts in the indicated anatomical regions. Bar plot displaying differences in the proportions of seven fibroblast subsets in the indicated anatomical regions. **e** Bubble heatmap depicting the expression of representative genes of *IDO1*^+^ fibroblasts across different stromal cell subsets. **f** Radar plot displaying the pathway enrichment analysis results for the four fibroblast subsets in module 1. The colors in the circles reflect the *P-values*. **g** Representative immunofluorescence staining for CXCL8 (white), IDO1 (green), COL1A2 (red, a marker gene of fibroblasts), and the nuclear marker DAPI (blue) in HC-IT tissue and CRS-NP tissue. The white arrows indicate colocalization of IDO1, CXCL8, and COL1A2 in the NPs. Scale bar, 20 μm.

**Fig. 5:**
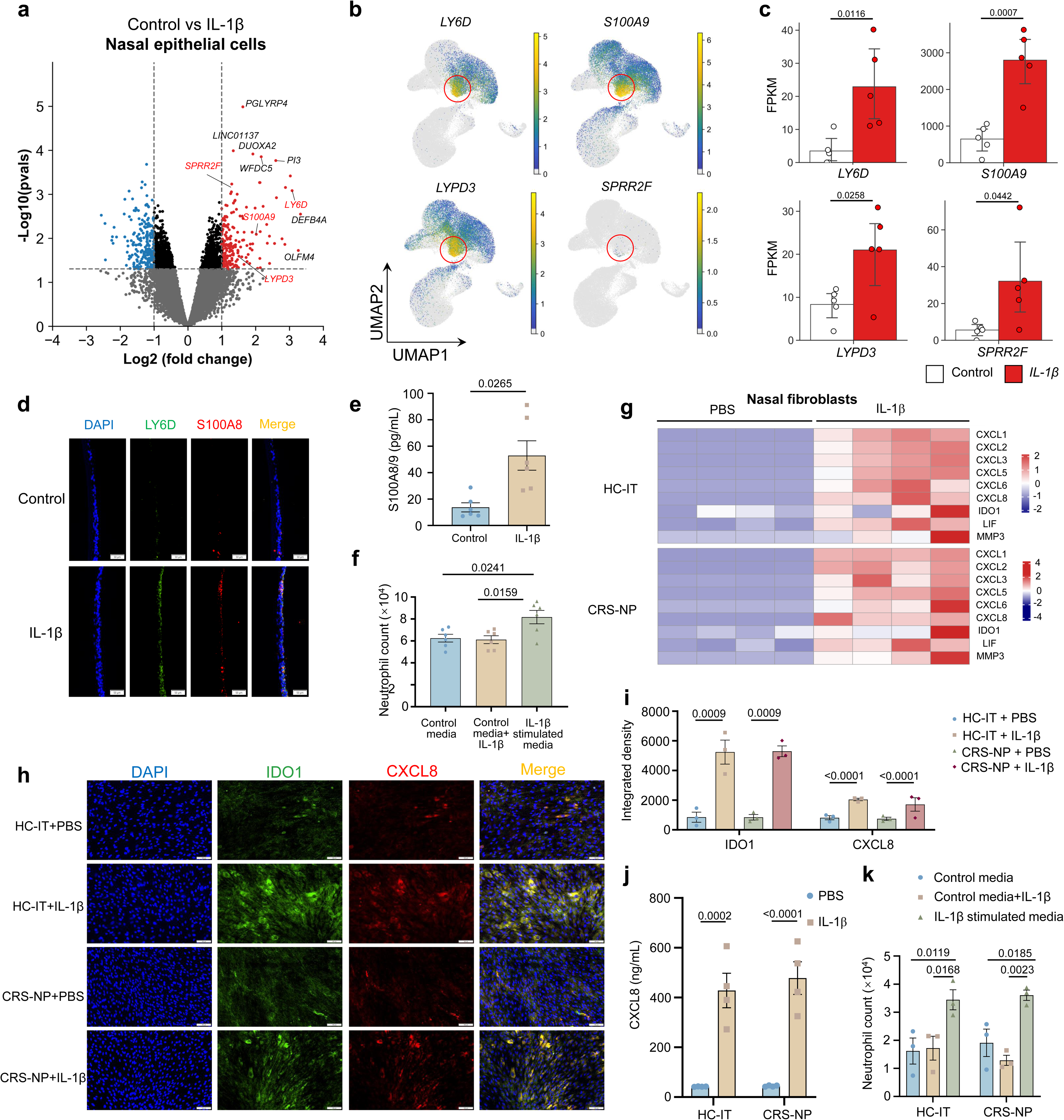
IL-1β induces transdifferentiation of *LY6D*^+^ club cells and *IDO1*^+^ fibroblasts to promote neutrophil recruitment. **a** Volcano plot of DEGs between IL-1β -stimulated and PBS-treated air-liquid interface (ALI) - cultured primary human nasal epithelial cells (HNEs) identified with the cut-off criterion *P* < 0.05 and |log2FC| ≥ 1. The *P-value*s were calculated and reported using two-tailed Student’s t-tests. Blue dots: significantly downregulated genes; red dots: significantly upregulated genes. **b** UMAP with epithelial cell compartment displaying the expression of four genes that are upregulated in *LY6D*^+^ club cells identified by scRNA-seq. The red circles indicate *LY6D*^+^ club cells in the epithelial cell compartment according to the scRNA-seq data. **c** Expression of four genes in panel b of ALI-cultured primary HNEs treated with the indicated conditions (as determined by bulk RNA-seq). The length of the error bars is a 95% confidence interval for the mean in Fig. 5(c). The *P-value*s were calculated and reported using a two-tailed Student’s t-test. **d** Representative immunofluorescence staining for LY6D (green), S100A8 (red), and the nuclear marker DAPI (blue) in ALI-cultured primary HNEs upon the indicated stimulations. Scale bar, 50 μm. **e** S100A8/A9 protein levels in the media (measured by ELISA) upon the indicated stimulations of ALI-cultured HNEs. The *P-value* was calculated and reported using a two-tailed Student’s t-test. **f** Number of neutrophils passing through the membrane of a transwell insert by ALI-cultured HNE-exposed media in the presence or absence of IL-1 stimulation. ALI-cultured HNE-exposed media (control media) and fresh media containing IL-1β (control media + IL-1β) were used as negative control. Data are presented as the means ±SEMs. The *P-value*s were calculated and reported using one-way ANOVA. **g** Heatmap displaying the expression of the indicated chemokines in primary fibroblasts derived from HC-IT tissue and CRS-NP tissue upon the indicated stimulations. **h** Representative immunofluorescence staining for IDO1 (green), CXCL8 (red) and the nuclear marker DAPI (blue) displaying the protein levels of IDO1 and CXCL8 upon the indicated stimulations. Scale bar, 100 μm. **i** Quantification of the data in panel (**h**). The data are presented as the means ± SEM. The *P-value*s were calculated and reported using a one-way ANOVA. **j** CXCL8 protein levels measured by ELISA after indicated stimulations in primary fibroblasts derived from HC-IT tissues and CRS-NP tissues. *The P-value*s are calculated and reported using two-way ANOVA. **k** Number of neutrophils passing through the membrane of a transwell insert by fibroblasts-exposed media in the presence or absence of IL-1β stimulation. Fibroblast-exposed media (control media) and fresh media containing IL-1β (control media + IL-1β) were used as negative controls. The data are presented as the means ± SEM. The *P-value*s were calculated and reported using two-way ANOVA.

Pathway enrichment analysis revealed that the transcriptome of *LY6D*^+^ club cells was enriched in genes induced by IL-1 signaling (Fig. 3i). The RNA velocity profile of total club cells indicated that *LY6D*^+^ club cells originated from resident club cells, suggesting that some club cells in the face of upregulated IL-1 signaling progressively acquired *LY6D*^+^ club cell identity in the mucosal epithelium in CRSwNP patients (Fig. 3j). The expression of several key functional genes and transcription factors upregulated during the maturation process of *LY6D*^+^ club cells was presented in the heatmap (Fig. 3k). IL1RN was inferred by RNA velocity, iteratively indicating that IL-1 signaling participates in the transdifferentiation of *LY6D*^+^ club cells (Fig. 3k). Pathway enrichment analysis revealed that genes involved in neutrophil degranulation were also enriched in *LY6D*^+^ club cells, reflecting the regulation of neutrophil recruitment by *LY6D*^+^ club cells (Fig. 3i). Taken together, these findings underscore the role of *LY6D*^+^ club cells in IL-1 signaling-mediated neutrophilic inflammation in CRSwNP.

### *IDO1*^+^ fibroblasts secrete chemokines that facilitate neutrophil recruitment in CRSwNP

To identify the stromal cell subsets responsible for inflammation in CRSwNP, we further sub-clustered the stromal cell compartment. Endothelial cells, pericytes and smooth muscle cells did not show much variation in the proportions of cell subsets across different anatomical regions except for an increased proportion of arterial endothelial cells and decreased proportion of lymphatic endothelial cells in CRS-related tissues as compared to those in HC-ITs, suggesting weakened lymphatic infiltration but enhanced angiogenesis in inflammatory tissues (Fig. 4a and Extended Data Fig. 7a-d). Fibroblast subsets exhibited substantial disparities in cellular proportions within the stromal cell compartment. The proportions of *IDO1*^+^ and *OXTR^+^* fibroblasts were markedly higher in NPs than in other tissues (Fig. 4a).

To better characterize the functionality of the fibroblast subsets, we proceeded to construct a transcription factor fate decision tree for different cell subsets within the stromal cell compartment. Six out of the seven distinct fibroblast subsets were categorized into two main modules (Fig. 4b, c). We noticed that module 1 comprised fibroblasts that exerted pro-inflammatory effects by enhanced production of certain chemokines, such as CXCL1 and CXCL8, while module 2 encompassed fibroblasts that mainly reside in the adventitia and are essential for regulating the integrity and function of the vessel structure^29–31^. The fibroblast clusters were displayed according to different anatomical regions (Fig. 4d). Within the cell clusters in module 1, *IDO1*^+^ and *OXTR*^+^ fibroblasts were enriched in inflammatory tissues, mostly in NPs, and they were barely detected in healthy tissues (Fig. 4d).

Considering the potent interaction detected between *IDO*1^+^ fibroblasts and neutrophils, we explored the gene expression patterns of different stromal cell subsets. The transcriptome of *IDO1*^+^ fibroblasts was enriched in chemokines (CXCL1/2/3/8) that are relevant to neutrophilic inflammation (Fig. 4e). We subsequently conducted pathway enrichment analysis on the fibroblast subsets within module 1, whose gene expression pattern was associated with inflammatory responses, to scrutinize the regulatory pathways in which *IDO1*^+^ fibroblasts are implicated (Fig. 4f). Interestingly, both the IL-1 signaling pathway and the NF-κB signaling were enriched in *IDO1*^+^ fibroblasts, indicating the upstream regulation of *IDO1*^+^ fibroblasts by IL-1 signaling in inflammation development. The high expression of *MMP3* and *LIF* in *IDO1*^+^ fibroblasts also indicated the regulation of *IDO1*^+^ fibroblasts by IL-1 signaling^32,33^ (Fig. 4e). Immunofluorescence staining revealed increased CXCL8 protein level and a greater number of IDO1^+^ fibroblasts (IDO1^+^ COL1A2^+^ cells) in CRS-NP samples as compared to HC-IT samples (Fig. 4g). Together, these findings elucidate IL-1 signaling as a common pathway inducing transdifferentiation of *LY6D*^+^ club cells and *IDO1*^+^ fibroblasts to facilitate neutrophil recruitment in CRSwNP.

### IL-1β−induced the transdifferentiation of *LY6D*^+^ club cells and *IDO1^+^* fibroblasts promotes neutrophil recruitment

IL-1 signaling can be activated by the interaction between IL-1β and IL-1 receptor (IL-1R), leading to various immune responses including neutrophilic inflammation^34^. Here we deployed recombinant IL-1β on air-liquid interface (ALI) cultures generated from primary human nasal epithelial cells (HNEs) (Extended Data Fig. 8a). Bulk RNA-seq data revealed that the addition of IL-1β elicited the *LY6D*^+^ club cell state of ALI-cultured HNEs, as IL-1β-stimulated ALI-cultured HNEs highly expressed genes that were also upregulated in *LY6D*^+^ club cells detected by scRNA-seq, such as *LY6D*, *SPRR2F*, *S100A9*, and *LYPD3* (Fig. 5a-c). Immunofluorescence staining of ALI-cultured HNEs revealed the colocalization and elevation of LY6D and S100A8 upon IL-1β stimulation (Fig. 5d). These results suggested that IL-1β induced transdifferentiation of *LY6D*^+^ cells *in vitro*. ELISA detected the increased secretion of S100A8/A9 protein from ALI-cultured HNEs upon IL-1β stimulation (Fig. 5e). Considering the ability of S100A8, S100A9, and S100A8/A9 to promote neutrophil activation, chemotaxis and adhesion, we performed a chemotaxis assay to determine whether the secretion of S100A8/A9 contributes to IL-1β-mediated neutrophil recruitment^26^. The results demonstrated that the media of ALI-cultured HNEs stimulated with IL-1β possessed a stronger neutrophil chemotactic capacity than the control media (Fig. 5f and Extended Data Fig. 8b).

To explore the effect of IL-1β on the induction of fibroblasts, we treated cultured primary fibroblasts isolated from IT tissues and NPs with IL-1β and performed bulk RNA sequencing. Bulk RNA sequencing results revealed high expression of genes encoding neutrophil chemoattractants (*CXCL1, CXCL2, CXCL3, CXCL5, CXCL6,* and *CXCL8*) in fibroblasts upon IL-1β stimulation (Fig. 5g). These genes were also upregulated in *IDO1*^+^ fibroblasts according to scRNA-seq analysis, suggesting that the primary fibroblasts acquire the identity of *IDO1*^+^ fibroblasts upon IL-1β stimulation. Immunofluorescence staining demonstrated that IL-1β was capable of activating fibroblasts and inducing the expression of CXCL8 and IDO1 in both IT-derived and NP-derived fibroblasts (Fig. 5h, i). In line with the results of bulk RNA sequencing, ELISA showed an increase in CXCL8 secretion in fibroblasts treated with IL-1β compared to those treated with PBS (Fig. 5j). As expected, culture media from IL-1β-exposed human nasal primary fibroblasts resulted in an increase in the transmigration of purified blood neutrophils compared with media of normal fibroblasts or fresh media mixed with IL-1β (Fig. 5k). Therefore, we reason that IL-1β induces both epithelial cells and fibroblasts to promote the recruitment of neutrophils to the sites of inflammation in CRSwNP.

### IL-1β antagonist impedes the transdifferentiation of *LY6D*^+^ club cells and *IDO1^+^* fibroblasts and mitigates inflammation *in vivo*

IL-1β is associated with neutrophilic airway inflammation^35^. Here we revealed increased IL-1β level in CRS-NP compared with that in HC-IT (Fig. 6a), most of which was expressed in MNP/DCs (Fig. 6b-d). The proportion of monocytes was greater in NP tissues than in other tissues, explaining an increase in IL-1β in the inflammatory mucosa (Fig. 6e). IL-1β is correlated with neutrophilic inflammation in CRS, which is frequently associated with worse disease outcomes^6^. However, whether therapy targeting IL-1β mitigates neutrophilic inflammation in CRSwNP is unknown.

**Fig. 6:**
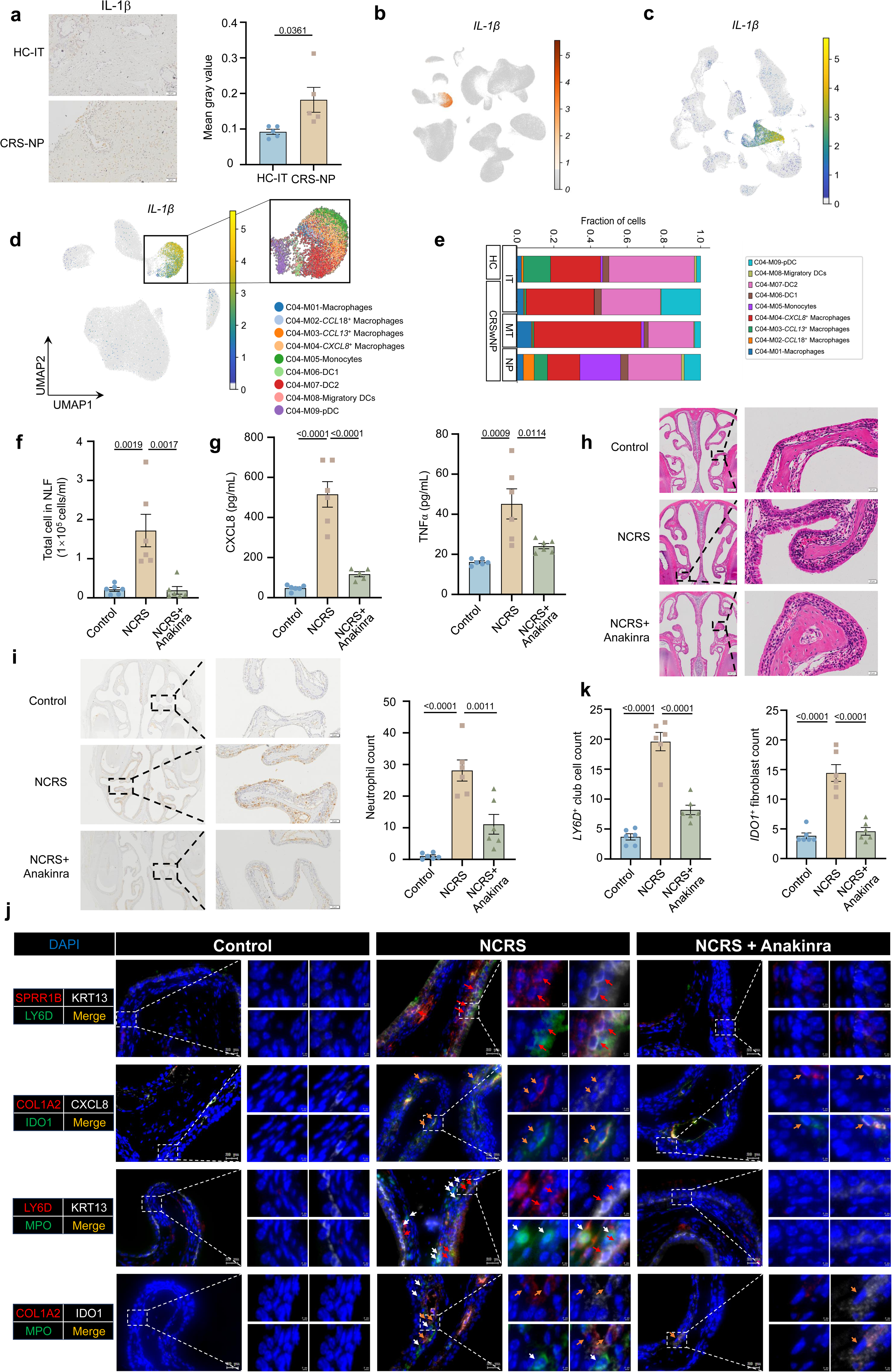
IL-1β antagonist suppresses transdifferentiation of *LY6D*^+^ club cells and *IDO1*^+^ fibroblasts and mitigates inflammation *in vivo*. **a** Representative immunohistochemistry staining for IL-1β in HC-IT tissue and CRS-NP tissue (n=5). Scale bar, 50 μm. The data are presented as the means ± SEM. The *P-value* was calculated and reported using a two-tailed Student’s t-test. **b** UMAP displaying the expression of IL-1β in total cell subsets determined via the scRNA-seq analysis. **c** UMAP showing the expression of IL-1β in total cell subsets from analysis of the CRSwNP scRNA-seq dataset (HRA000772). **d** UMAP embedding the expression of IL-1β in immune cells determined via the scRNA-seq analysis. **e** Bar plot depicting the cell compositions of MNP/DC subsets for the indicated anatomical regions of human nasal mucosa from CRSwNP patients and healthy individuals. **f** Cell counts in nasal lavage fluid from mice in the indicated groups. Data are expressed as the means ± SEM. The *P-value*s were calculated and reported using one-way ANOVA. **g** CXCL8 (left) and TNFα (right) protein levels in the nasal lavage fluid of mice in the indicated groups measured by ELISA. The data are expressed as the means ± SEM. The *P-value*s were calculated and reported using a one-way ANOVA. **h** Representative H&E images of nasal mucosal tissues from mice in the indicated groups. Scale bar, 200 μm (left), 20 μm (right). **i** Representative immunohistochemical staining for MPO in nasal mucosal tissues of mice from the indicated groups (left). Scale bar, 200 μm (left), 50 μm (right). The data are presented as the means ± SEM (right). The *P-value*s were calculated and reported using one-way ANOVA. **j** Representative multiple immunohistochemistry images of nasal mucosa in mice from the indicated groups. Images showing the the staining for *IDO1*^+^ fibroblasts (orange arrows), *LY6D*^+^ club cells (red arrows), and neutrophils (white arrows) in the nasal mucosa of model mice. Scale bar, 20 μm (left), 2 μm (right). **k** Quantification of (**j**). Data are presented as the means ± SEM. One-way ANOVA was employed to assess variations of *IDO1*^+^ fibroblasts and *LY6D*^+^ club cells.

To explore the effect of IL-1β inhibition on neutrophilic CRSwNP, we established a mouse model of lipopolysaccharide (LPS)-induced neutrophilic chronic rhinosinusitis with nasal poly (NCRS)^36^, and then treated the model mice with anakinra, a recombinant, nonglycosylated interleukin-1 receptor antagonist that has been employed as a therapeutic intervention for autoinflammatory diseases and hematological malignancies^37,38^ (Extended Data Fig. 8c). The total cell count in nasal lavage fluid (NLF) from mice serves as an indicator of inflammation severity^39^. We detected fewer cells in the NLF from anakinra-treated NCRS mice as compared to untreated NCRS mice (Fig. 6f). ELISA detected elevated secretion of CXCL8 and TNFα in the NLF from NCRS mice, while the secretion of these factors approached to normal level in anakinra-treated NCRS mice (Fig. 6g). In NCRS mice, we observed the inflammatory features represented by increased inflammatory cell infiltration and mucosal hyperplasia with impaired mucosal integrity, which were alleviated in anakinra-treated NCRS mice (Fig. 6h). Immunochemistry staining detected increased neutrophil infiltration in NCRS mice, which was also improved in anakinra-treated NCRS mice (Fig. 6i). These findings reflected the substantial mitigation of inflammation by IL-1β blockade in NCRS mice.

Similar to the induction of human primary cells by IL-1β, we detected increased numbers of *LY6D*^+^ club cells and *IDO1*^+^ fibroblasts in the mucosa of NCRS mice as compared to those in control mice (Fig. 6j, k). We next investigated whether IL-1β antagonist affects the transdifferentiation of *LY6D*^+^ club cells and *IDO1*^+^ fibroblasts in NCRS mice. Immunofluorescence staining demonstrated that numbers of LY6D^+^ club cells and IDO1^+^ fibroblasts were declined in the mucosa of anakinra-treated NCRS mice, along with the reduction of neutrophil infiltration, as compared to untreated NCRS mice (Fig. 6j, k). These findings suggested that IL-1β suppression impedes the transdifferention of *LY6D*^+^ club cells and *IDO1*^+^ fibroblasts and mitigates neutrophilic inflammation, suggesting that targeting IL-1β is an effective intervention against neutrophil recruitment in CRSwNP.

## Discussion

Here, we presented a detailed profile of the nasal mucosa of HC-ITs, CRS-ITs, CRS-MTs, and NPs from CRSwNP patients at the single-cell level. We identified *LY6D*^+^ club cells and *IDO1*^+^ fibroblasts in the nasal mucosa that promote neutrophil recruitment in CRSwNP. *LY6D*^+^ club cells exert the pathogenic effects upon IL-1 signaling stimulation by secreting S100A8 and S100A9, two molecules possessing ability to promote neutrophil chemotaxis^26^. In addition, *IDO1*^+^ fibroblasts induced by IL-1 signaling produce multiple chemokines that interact with receptors expressed in neutrophils and promote neutrophil recruitment. IL-1β, a key factor in the IL-1 signaling pathway, was demonstrated to be upregulated in NPs from CRSwNP patients. We found that IL-1β induces the transdifferentiation of *LY6D*^+^ club cells and *IDO1*^+^ fibroblasts from epithelial cells and fibroblasts, respectively. Increased numbers of *LY6D*^+^ club cells and *IDO1*^+^ fibroblasts were also observed in NCRS mouse model. Administration of an IL-1β antagonist reduced the numbers of *LY6D*^+^ club cells and *IDO1*^+^ fibroblasts, and showed a promising effect on alleviating neutrophilic inflammation in NCRS mice (see the model in Extended Data Fig. 8d).

A previous study detected higher mRNA and protein levels of IL-1β in NPs than in uncinate tissues, inferior turbinates, and ethmoid sinus mucosal samples from control subjects, as did an increased number of IL-1β^+^ cells in polyp tissue from neutrophilic CRSwNP patients^40,41^. However, the cell sources of IL-1β in the nasal mucosa are unclear. Here, we verified the upregulation of IL-1β in NP samples from CRSwNP patients and identified that the cell sources of IL-1β in CRSwNP were monocytes, macrophages, DCs, and neutrophils. Our results elucidated the role of IL-1β in determining the transdifferentiation of *LY6D*^+^ club cells and and *IDO1^+^*fibroblasts, both of which are enriched in NP tissues. Elevated expression of *S100A8* and *S100A9* has been observed in nasal polyps compared to control tissues^9,23,24^. Both proteins that induce neutrophil chemotaxis and adhesion^26^ are secreted by *LY6D*^+^ club cells in the epithelium from the nasal mucosa. Elevated levels of EGFR ligands have been detected in various airway disorders, such as CRS and COPD^42,43^. Our results also demonstrated increased expression of *EREG* and *AREG* in *LY6D*^+^ club cells, indicating the involvement of these cells in activating EGFR signaling and subsequently inducing mucus and inflammatory cytokine secretion from airway epithelial cells^42,44^. The functionality of *LY6D^+^* club cells is multifaceted and deserves further exploration. Previous studies have shown that immune cells and stromal cells within the organs, including macrophages and fibroblasts, send coordinated signals that guide neutrophils to their final destination^45,46^. Our data uncovered an unreported mechanism underpinning neutrophil chemotaxis orchestrated by *IDO1*^+^ fibroblasts in CRSwNP. *IDO1*^+^ fibroblasts constitute the core cell subset that promotes neutrophil recruitment based on the strong interaction observed between these two cell subsets in CRSwNP. IL-1β-induced *IDO1*^+^ fibroblasts release substantial quantities of chemokines (CXCL1/2/3/5/6/8) to promote neutrophil recruitment. Considering that both *LY6D^+^* club cells and *IDO1*^+^ fibroblasts are induced by IL-1β, future studies should examine whether other pro-inflammatory cytokines in the IL-1 signaling pathway, such as IL-1α, contribute to the transdifferentiation of the two cell subsets.

It is well known that neutrophilia and eosinophilia are both present in most cases of CRS^19^. Activated neutrophils possess the capability to facilitate eosinophil transmigration and accumulation^47^. Studies have demonstrated the association of mixed eosinophilic-neutrophilic inflammation with hard-to-treat asthma or CRSwNP^5,19,48^. In addition to surgery and intranasal corticosteroids, multiple biologics have been approved or are undergoing clinical trials as therapeutics for CRS. Dupilumab (targeting IL-4Rα), omalizumab (targeting IgE), and mepolizumab (targeting IL-5) have been approved for CRSwNP treatment. Reslizumab (targeting IL-5) and benralizumab (targeting IL-5Rα) have been undergoing phase 2 and phase 3 trials, respectively^49^. However, these therapies primarily target eosinophilic and type 2 inflammation in CRSwNP, whereas therapies targeting neutrophilic inflammation remain a gap. Given the unfavorable prognosis of CRSwNP with a mixed inflammatory pattern and the ineffectiveness of steroids on the neutrophil activation state in CRSwNP, the demand to develop novel strategies against neutrophilia in CRSwNP patients is imperative^49^. Strategies targeting IL-1β, such as anakinra, rilonacept, and canakinumab, are commonly used to block the effects of IL-1β, thereby reducing inflammation and related symptoms in conditions such as rheumatoid arthritis, atherosclerosis, and other immune-mediated diseases^50^. In this study, we revealed the effects of intervention targeting IL-1β on inducing *LY6D*^+^ club cells and *IDO1*^+^ fibroblasts, and recruiting neutrophils in CRSwNP. We validated the mitigation of neutrophilic inflammation by the application of anakinra in an LPS-induced neutrophilic CRSwNP mouse model. We did observe a substantial reduction in chemokine secretion and a decrease in neutrophil infiltration in anakinra-treated neutrophilic CRSwNP mice. Given the promising anti-inflammatory effects of blocking IL-1β in the neutrophilic CRSwNP mouse model, our findings highlight that targeting IL-1β may be an effective strategy for the treatment of neutrophilic inflammation in CRSwNP.

## Materials and methods

### Study participants

In total, 85 individuals aged between 18 and 70 years were recruited from the Department of Otolaryngology in Qilu Hospital of Shandong University, including chronic rhinosinusitis with polyps (CRSwNP) patients (n=47) and healthy controls (HCs) (n=38). This study was approved by the Medical Ethics Committee of Qilu Hospital of Shandong University (KYLL-202102-1061). All study participants provided written informed consent. The diagnosis of CRSwNP was based on the EPOS 2020 criteria^51^, and included confirmatory clinical, endoscopic and radiographic criteria. HCs were patients with cerebral spinal fluid leak or nasal septum deviation. The nasal tissues, including nasal polyps, middle turbinates, and inferior turbinates, were collected during endoscopic sinus surgery. Participants who had an immunodeficiency disorder, fungal sinusitis, cystic fibrosis or tumors were excluded from the study. No participants used systemic corticosteroids for at least 4 weeks before surgery. The detailed clinical characteristics are summarized in Supplementary Table 1.

### Preparation of single-cell suspensions

Nasal mucosa was freshly sampled from the middle turbinates (n=7), inferior turbinates (n=9), nasal polyps (n=15) of CRSwNP patients and inferior turbinates (n=2) of patients with cerebral spinal fluid leak. The nasal biopsies were washed in phosphate-buffered saline (PBS, 10010023, ThermoFisher) to remove mucus and blood cells. Then, the nasal tissues were cut into approximately 0.5-mm^3^ pieces in RPMI-1640 medium supplemented with 1% penicillin/streptomycin, and then enzymatically digested with the Multi Tissue Dissociation Kit 2 (MACS# 130-110-203) at 37℃ for 30 min with agitation, according to the manufacturer’s instructions. Following cell dissociation, the resultant cell suspension was sequentially filtered through cell strainers with pore sizes of 70 μm and 40 µm (BD). Subsequently, the samples were centrifuged at 300g for 10 minutes. Subsequent to the removal of the supernatant, the cells forming the pellet were reconstituted in red blood cell lysis buffer (Thermo Fisher) and subjected to a 2-minute incubation on ice to lyse the red blood cells. Following dual washes with PBS, the cellular pellets were re-suspended in PBS supplemented with 0.04% bovine serum albumin (A7906, Sigma-Aldrich).

### Single-cell RNA library construction and sequencing

DNBelab C Series High-throughput Single-cell System (BGI-research) was utilized for scRNA-seq library preparation. Briefly, the single-cell suspensions underwent a series of processes to generate barcoded scRNA-seq libraries. These steps encompassed droplet encapsulation, emulsion breakage, collection of beads containing the captured mRNA, reverse transcription cDNA amplification and subsequent purification. The cDNA was subjected to fragmentation into shorter segments spanning 250 to 400 base pairs. Following this, the construction of indexed sequencing libraries was achieved in accordance with the protocol provided by the manufacturer. Qualification was performed using the Qubit ssDNA Assay Kit (Thermo Fisher Scientific) and the Agilent Bioanalyzer 2100. Subsequent to library preparation, all the constructs underwent sequencing using the DIPSEQ T1 sequencing platform in the China National GeneBank via pair-end sequencing methodology. The sequencing reads contained 30-bp read 1 (including the 10-bp cell barcode 1, 10-bp cell barcode 2 and 10-bp unique molecular identifiers [UMI]), 100-bp read 2 for gene sequences and 10-bp barcodes read for sample index. Next, processed reads were aligned to the GRCh38 reference genome using STAR (v2.5.3). The identification of valid cells was achieved through an automated process utilizing the “barcodeRanks” function from the DropletUtils tool. This function was employed to eliminate background beads and those with UMI counts falling below a predetermined threshold, using the UMI number distribution characteristic of each cell. Finally, we computed the gene expression profiles of individual cells and subsequently generated a matrix of genes by cells for each library by means of PISA. The newly generated scRNA-seq data and bulk RNA-seq data will be immediately accessible upon acceptance of the paper.

### Alignment, quantification, and quality control of single-cell RNA sequencing data

The droplet-based sequencing data were subjected to alignment and quantification through the utilization of CellRanger software (version 3.0.2, designed for 3’ chemistry), employing the GRCh38.p13 human reference genome. The Python package Scanpy (version 1.7.1)^52^ was employed to load the matrix containing cell-gene counts and to execute quality control procedures for both the newly generated dataset and the acquired datasets. For each sample, genes associated with mitochondria (indicated by gene symbols commencing with “MT-”) and ribosomal proteins (initiated by gene symbols commencing with “RP”) were eliminated from consideration. After that, cells possessing less than 2000 UMI counts and 250 detected genes were identified as empty droplets and subsequently excluded from the datasets. Finally, genes demonstrating expression in fewer than three cells were excluded from further analysis.

### Doublet detection

In order to rule out doublets, we implemented the Scrublet software (version 0.2.3)^53^, which facilitated the identification of artifactual libraries originating from two or more cells within each scRNA-seq sample, comprising both the newly generated dataset and the compiled datasets. The doublet score for each individual single cell, along with the threshold determined from the bimodal distribution, was computed using the default parameters (sim_doublet_ratio=2.0; n_neighbors=None; expected_doublet_rate=0.1, stdev_doublet_rate=0.02). After that, a comprehensive assessment was conducted on the remaining cells and cell subsets to identify potential false-negatives from the scrublet analysis. This evaluation was guided by the following sets of criteria: (1) cells with more than 8000 detected genes, (2) subsets that expressed marker genes from two distinct cell types, which are unlikely according to prior knowledge (i.e., CD3D for T cells and EPCAM for epithelial cells). Any cells or subsets identified as doublets were excluded from subsequent downstream analyses.

### Graph subseting and partitioning cells into distinct compartments

Downstream analysis included normalization (scanpy.pp.normalize_total method, target_sum=1e4), log-transformation (scanpy.pp.log1p method, default parameters), cell cycle score (scanpy.tl.score_genes_cell_cycle method), cell cycle genes defined in Tirosh et al, 2016^54^, feature regress out (scanpy.pp.regress_out method, UMI counts, percentage of mitochondrial genes and cell cycle score were considered to be the source of unwanted variability and were regressed), feature scaling (scanpy.pp.scale method, max_value=10, zero_center=False), PCA (scanpy.tl.pca method, svd_solver=’arpack’), batch-balanced neighborhood graph building (scanpy.external.pp.bbknn method, n_pcs=20)^55^, leiden graph-based subseting (scanpy.tl.leiden method, Resolution=1.0)^56^, and UMAP visualization^57^ (scanpy.tl.umap method) performed using scanpy. The initial categorization of the subsets encompassed a division into six distinct compartments, achieved through the utilization of marker genes established in the existing literature in conjunction with genes exhibiting differential expression. (scanpy.tl.rank_gene_groups method, method=’Wilcoxon test’). Specifically, the epithelial compartment was annotated using a gene list (EPCAM, KRT8, KRT18, KRT19, PIGR), T and ILCs compartment (CD2, CD3D, CD3E, CD3G, TRAC, IL7R), B cell compartment (JCHAIN, CD79A, IGHA1, IGHA2, MZB1, SSR4), MNPs compartment (HLA-DRA, CST3, HLA-DPB1, CD74, HLA-DPA1, AIF1), Mast cell compartment (TPSAB1, CPA3, TPSB2, CD9, HPGDS, KIT), and Stromal cell compartment (IGFBP7, IFITM3, TCF7L1, COL1A2, COL3A1, GSN). Subsequently, the epithelial compartment was subjected to sorting for subsequent downstream analysis.

### Transcription factor module analysis

The python package pySCENIC workflow (version 0.11.0) with default settings was used to infer active TFs and their target genes in all human cells^58,59^. Specifically, the pipeline was executed in three steps. Initially, the single-cell gene expression matrix was filtered to eliminate genes whose expression was detected in fewer than ten total cells. The retained genes were subsequently employed to construct a gene-gene correlation matrix, which facilitated the identification of co-expression modules through the application of a regression per-target approach utilizing the GRNBoost2 algorithm. Subsequent to the initial step, each identified module was systematically refined based on a regulatory motif in close proximity to a transcription start site (TSS). The acquisition of cis-regulatory footprints was facilitated through the utilization of positional sequencing methodologies. The binding motifs of the TFs were then used to build an RCisTarget database. Modules were retained based on the enrichment of transcription factor (TF)-binding motifs among their respective target genes. In cases where target genes lacked direct TF-binding motifs, they were excluded from consideration. In the third phase, we assessed the influence of each regulon on individual single-cell transcriptomes through the utilization of the area under the curve (AUC) score, employing the AUCell algorithm as the evaluative metric. The scores pertaining to transcription factor motifs within gene promoters and regions surrounding transcription start sites, specific to the hg38 human reference genome, were acquired from the RcisTarget database. Concurrently, the list of transcription factor-associated genes was obtained from the Humantfs database^60^.

### Fate decision tree construction (regulon-based)

Dendrogram plots were generated for epithelial cells using the sc.pl.dendrogram method from the Scanpy package. These plots were generated based on the AUCell matrix comprising 608 regulons, aiming to visualize more nuanced alterations. We deciphered the diverging composite rules of a regulon-based dendrogram by testing each branching node for differential regulon importance. Thereafter, differential analysis of regulon expression was conducted for each node using the Wilcoxon test (implemented through the sc.tl.rank_gene_groups method with method=’Wilcoxon test’), with the aim of deducing the sequence of regulon-driven propagation events.

### Datasets integration

In this study, we utilized a previously published scRNA-seq dataset of CRSwNP^20^ (GSA: HRA000772), the one that detailed a specific quantity of neutrophils, to investigate the expression of inflammatory factors in neutrophils in human nasal mucosal tissues. Specifically, we compared the downloaded fastq files with the barcodes-genes matrix utilizing Alevin-fry^61^. The matrix underwent initial quality control, doublet removal, and normalization, applied in accordance with the dataset from the previous section. The gene expression and cell annotation of the dataset were modeled using CellTypist^62^. Subsequently, the trained model was used to perform Label Transfer on the HRA000772 dataset. In particular, myeloid cells annotated by Label Transfer were manually reannotated based on marker genes, thereby identifying the neutrophils subset (*FCGR3B^+^CXCR1^+^CXCR2^+^*).

### RNA velocity

Cells that met the quality control criteria were used to filter the loom file generated by the Velocyto python package based on the cell barcodes^63^. This package was used to conduct splicing analysis on the bam file in preparation for subsequent RNA velocity analysis. The filtered loom file served as an input within the Scanpy pipeline, implemented as part of the CellRank pipeline^64^. The loom file derived from Velocyto was harnessed to compute RNA velocities for each cell according to standard parameters for the software. CellRank generates both stochastic and dynamic models of RNA velocity, which were compared via the computation of a consistency score for each cell, employing each modeling approach, in accordance with the guidance provided by the authors. Pseudotime was subsequently calculated based on the outcomes of RNA velocity analysis, while latent time was deduced from the dynamic velocity results.

### Gene set scoring and identification of significant changes

We scored the gene sets of all cells and subsets using the Scanpy python package (sc.tl.score_genes method, ctrl_size=len(genesets), gene_pool=None, n_bins=25, use_raw=None). The score was the average expression of a set of genes subtracted from the average expression of a reference set of genes. The reference set was randomly sampled from the gene_pool for each binned expression value. To prevent highly expressed genes from dominating a gene set score, we scaled each gene of the log2 (TP10K+1) expression matrix by its root mean squared expression across all cells. After obtaining score-cell matrix of the signatures, differential signature analysis (sc.tl.rank_gene_groups method, method=’Wilcoxon test’) was implemented to identify significant changes among different nasal anatomical regions. All pathways included in gene set enrichment analysis (Fig. 3i, Fig. 4f and Extended Data Fig. 7c, d) were obtained from Reactome^65^.

### Cell-cell interaction and network representation analysis

To plot chemokine-chemokine receptor interaction networks, we employed the Networkx (version 2.5) (https://github.com/networkx/networkx), Community (version 1.0.0b1) and Pygraphviz (version 1.6) (https://github.com/pygraphviz/pygraphviz) python packages to construct a network defined using the count of interactions between cell subsets. The pipeline was implemented in three steps. First, the nodes with a degree of zero were eliminated. Second, any edges with a connection strength less than the average of all the edges were removed. Third, the sizes of the nodes were defined as the log2 (counts+1) of the cell subsets, and the network with the Kamada Kawai layout algorithm (networkx.kamada_kawai_layout method) was utilized to visualize the network. The thickness of the line connecting the two cell subsets was directly proportional to the degree of interaction strength between them. The chemokines-chemokines receptor interaction data were obtained from IMEx Consortium^66^, IntAct^67^, InnateDB-All^68^, MINT^69^ and I2D^70^ database.

### Estimation of neutrophil infiltration in CRSwNP

In this study, we applied the xCell algorithm to determine the immune cell subsets in the RNA-seq dataset (GSE179265). The xCell algorithm represents a gene signature-based approach derived from learning from numerous pure cell types originating from diverse sources. This method adeptly enables a cell type enumeration analysis using gene expression data, providing a comprehensive assessment of 64 immune and stromal cell types. This attribute endows it with a commendable capability to accurately depict the intricate landscape of cellular heterogeneity within tissue expression profiles^71^.

### Animals

C57BL/6 mice used in these experiments were purchased from SPF Biotech. The mice were maintained in individually ventilated cages in a specific pathogen-free facility under 12 h light–dark cycles at 22–24 °C and 50–60% humidity. The protocol for the animal studies was approved by the Laboratory Animal Ethical and Welfare Committee of Shandong University Cheeloo College of Medicine (23086).

### Neutrophilic CRSwNP mouse model and treatment with an IL-1R antagonist (Anakinra)

Mice were randomly divided into three groups consisting of 6 individuals each. The construction of the mouse model of CRSwNP with neutrophilia was carried out following a previously described protocol^36^. For the control group, 20 µl of sterile normal saline solution was dropped into the nasal cavities three times a week for 3.5 consecutive months. Mice in the model groups received 10 µg of LPS (from *Escherichia coli*; Sigma-Aldrich, Merck Millipore, Germany) in 20 µl of sterile normal saline solution three times a week for 3.5 consecutive months. For the anakinra-treated group, starting on the 77th day, the mice were given 10 µg of Anakinra (MedChemExpress, HY-108841, USA) in 20 µl of sterile normal saline solution by intranasal instillation and 10 µg of Anakinra in 200 µl of saline by intraperitoneal injection 30 minutes after LPS stimulation for 2 weeks. For the following 2 weeks, only 10 µg of Anakinra was intranasally administered in 20 µl of sterile normal saline solution within 30 minutes each after LPS stimulation. The animals were sacrificed 24 h after the last nasal challenge. The graphic protocol is depicted in Extended Data Fig. 8c. NLF was collected immediately from the sacrificed mice by washing the nasal cavity with 1 mL of ice-cold PBS three times. The total number of cells in NLF was counted using a cell counter (JIMBIO, China).

### Immunofluorescence staining

The detailed experimental protocol for processing the sinonasal tissue specimens was previously described^72^. In brief, we removed the skin on the heads of the mice and then excised the mandibles. The heads of the mice were fixed in 4% paraformaldehyde at room temperature for at least 24 hours, and decalcified for 7 days. For human nasal tissues, biopsy samples were soaked in 4% paraformaldehyde for 24 hours. For both the murine and human studies, after dehydration and paraffin embedding, the tissue samples were cut into 4 µm-thick paraffin sections. The slides were incubated at 65 ℃ for 1 hour, dewaxed, hydrated, and subsequently heated in antigen retrieval liquid for 15 minutes in a microwave oven. After cooling to room temperature, the slides were permeated with PBS containing 1% Triton X-100 for 20 minutes. The slides were washed in PBS 3 times and blocked with 5% bovine serum albumin at room temperature for 1 hour. After that, the slides were incubated with the primary antibody (see Supplementary Table 2 for a complete list and dilutions) overnight at 4℃ in a humidified chamber. The slides were gently washed with PBS 3 times, and incubated with a fluorescent secondary antibody at room temperature for 1 hour. After washing with PBS, the slides were stained with 4’, 6-diamidino-2-phenylindole ( DAPI) (Solarbio, C006, China) for 10 minutes. After another washing step with PBS, the slides were cover-slipped with anti-fade mounting medium (Solarbio, S2100, China). Image acquisition was performed using two fluorescence microscopes (Olympus, IX73 and VS120, Japan).

### Multiplexed immunohistochemistry

Multiplexed immunohistochemistry (mIHC) assay was performed using the Opal 6-Plex Detection Kit (AKOYA #811001, USA) as described previously^73^. Briefly, after dewaxing and hydration, the slides were boiled in AR6 buffer in a microwave oven for 15 minutes. The tissue sections on the slides were incubated with blocking buffer for 30 min and then with primary antibody (see Supplementary Table 2 for a complete list and dilutions) for 2 hours at room temperature in a humidified chamber. Then the slides were washed with TBST twice and incubated with Opal polymer anti-rabbit/mouse horseradish peroxidase (HRP) for 10 minutes at room temperature. Then, 100-300 µl of Opal Fluorophore working solution was added to each slide. After washing with TBST twice, the slides were incubated at room temperature for 10 minutes. The previous steps were repeated as needed. DAPI working solution was applied on the slides for 10 minutes at room temperature. As a final step, the slides were washed and cover-slipped with anti-fade mounting medium. Image acquisition was performed using the TissueFAXS imaging system (TissueGnostics, Germany).

### Isolation and culture of primary human nasal epithelial cells (HNEs)

Human nasal epithelial cells were scraped from patients’ nasal mucosa during endoscopic sinus surgery. The cells were placed in an Eppendorf tube containing 1 ml of bronchial epithelial cell medium (BEpiCM) (ScienCell, 3211, USA) supplemented with 1% penicillin/streptomycin and 1% bronchial epithelial cell growth supplement immediately upon acquisition. Cells were seeded within 6 hours in six well plates pre-coated with Collagen Type I (Corning, 354236, USA) and maintained in a humidified incubator at 37℃ containing 5% CO_2_. The media was changed every two days. When cells reached 90% confluence in the well, they were transferred to the upper chamber of polyester Transwell inserts (0.4 µm, 0.33 cm^2^, BIOFIL, TCS016012, China) pre-coated with Collagen Type Ⅰ. After that, 1 ml of BEpiCM was added into the lower chamber, and media was replaced every two days. At confluence, the media was replaced with differential media (BEpiCM: DMEM/F12 =1:1) in the basal chamber and the apical surface was exposed to provide an air-liquid interface (ALI). Monolayers were grown at the ALI for an additional 21 days to promote differentiation into a nasal epithelium with basal, multiciliated and secretory cells. On day 22, media containing PBS or recombinant IL-1β (10 ng/ml) (Abbkine, PRP100051, USA) was added to the basal chambers for 3 days.

### Isolation and culture of primary human nasal fibroblasts (HNFs)

The inferior turbinate or nasal polyp tissues were soaked in penicillin-streptomycin solution (Solarbio, P1400, China) for 3 minutes and cut into small pieces. After digestion in Trypsin-EDTA solution (Macgene, CC017-500) for 10 minutes, the tissues were put into cell culture flasks with DMEM media supplemented with 10% FBS. The cells were cultured in a humidified incubator at 37 ℃ containing 5% CO_2_, and the media was replaced every 2 days. The migrated cells were nasal mucosa-derived fibroblasts. When cells reached 90% confluence in the well, PBS or IL-1β (10 ng/ml) was added into the wells, and the cells were cultured for 1 days.

### Isolation of human peripheral blood neutrophils

Neutrophils were enriched from peripheral blood by means of Polymorphprep (Serumwerk Bernburg AG, 1895) density centrifugation. We carefully layered 5.0 ml of anti-coagulated whole blood over 5.0 ml of PolymorphPrep in a 15 ml tube. The tubes were centrifuged at 500 g for 30 min at 20℃. After centrifugation, two bands were visible, and the neutrophils were enriched in the lower band. The cells were aspirated to another clean tube and an equal volume of sterile normal saline solution was added. After incubating at room temperature for 10 minutes, the tubes were put on centrifuge at 500 × g for 30 minutes. The supernatant was discarded, and the cell pellet was resuspended in Roswell Park Memorial Institute (RPMI) 1640 media supplemented with 1% FBS.

### Neutrophil chemotaxis assay

For the cell migration assay, after resuspension in RPMI 1640 media supplemented with 1% FBS, the neutrophils were seeded at 1.0 × 10^5^/100 µl per well in the upper compartment of 24-transwell plates with 3-μm pores (Costar, 3415). The conditioned media from fibroblasts, either stimulated with IL-1β or not, was added into the lower chamber to test the chemotactic effect on neutrophils. Normal culture media was used as a negative control. After 3 hours of incubation at 37℃ in 5% CO_2_, the number of the migrated cells in the lower chamber was counted.

### Enzyme-linked immunosorbent assay (ELISA)

ELISAs were performed using multiple ELISA kits (4A Biotech, CHE0011, CME0008, CME0004, China) according to the manufacturers’ instructions. In brief, the standards and samples were added to the antibody pre-coated 96-well ELISA plate, which was subsequently incubated at 37℃ for 2 hours. The liquid was removed, and the plate was washed 4 times with wash buffer. Then, an enzyme-linked antibody was applied to the plate, which was incubated at 37℃ for 60 minutes. After a washing step, avidin-biotin-peroxidase complex was applied to each well, and the plate was incubated at 37℃ for 30 minutes. The plate was washed 4 times with wash buffer and the color developing reagent was added to each well of the plate and the plate was incubated at 37℃ in darkness for 10-20 minutes. The reaction was terminated by adding stop solution and the optical density (OD) at 450 nm was measured immediately using a microplate reader (Thermo Fisher, Varioskan Flash, USA). Analysis was performed using GraphPad Prism version 9.

### Immunohistochemistry

Paraffin-embedded sections were incubated at 65℃ for 1 hour. Dewaxing, hydration, and antigen repair were performed sequentially as previously described^73^. The endogenous peroxidase blocker was applied to the slides after they had cooled to room temperature. The slides were incubated for 20 minutes at room temperature. The slides were then washed with PBS 3 times and incubated with the primary antibody (see Supplementary Table 2 for a complete list and dilutions) in a humidified chamber at 4℃ overnight. After washing with PBS, the sections were incubated with reaction enhanced solution. Following another wash, the sections were incubated with the secondary antibody for 10 minutes, and the color reaction was developed using 3,30-diaminobenzidine tetrahydrochloride (DAB) (ZSGB-Bio, PV-9000, China). The slides were counterstained with hematoxylin. Finally, the slides were dehydrated and mounted. The images were acquired using a fluorescence microscope (Olympus VS120, Japan).

### Hematoxylin and eosin Staining (HE staining)

HE staining was performed using the HE staining kit (Beyotime, C0105S, China) according to the manufacturer’s instruction. Sections were dewaxed, hydrated and then washed with PBS. Then, the sections were incubated with hematoxylin for 10 seconds and washed with distilled water for 10 minutes. After that, the sections were differentiated with 1% hydrochloric ethanol for 20 seconds. After a washing step with distilled water for a 10 min, the slides were stained with eosin for 1 min. Following dehydration, clearing and mounting, the slides were ready for image acquisition under a microscope (Olympus, VS120, Japan).

### Statistical methods

No statistical analysis was performed to predetermine sample size. The numbers of samples included in the analyses are listed throughout the figures. For the scRNA-seq data, statistical analyses and graphic production were performed using Python version 3.7.10. The experimental data are presented as mean ± SEM or mean with 95% CI, as shown in the corresponding figure legends. Data distribution was assumed to be normal. One-way ANOVA and two-way ANOVA were used to compare multiple sets. Two-tailed Student’s t-tests were used for the comparisons between two sets. Statistical analyses and graphic production were performed with GraphPad Prism version 9 (GraphPad Software Inc., San Diego, CA, USA). *P* < 0.05 was considered statistically significant.

## ACKNOWLEDGMENTS

This research is supported by the National Natural Science Foundation of China (82171106, 82371120, 81700890), Taishan Scholar Program of Shandong Province (tsqn202103166), and Natural Science Foundation of Shandong Province (ZR2022MH313).

## AUTHOR CONTRIBUTIONS

X.F. and P.W. conceived and designed the research. X.X. and M.J. collected and processed the tissue to single-cell suspensions. Y.W. performed all computational analyses for scRNA-seq data. Y.W. and P.W. performed analyses for bulk mRNA sequencing data on epithelial cells and fibroblasts. X.X, Y.W., L.Q., S.G., and P.W analyzed data, prepared figures. X.X, C.W., Y.Y. and W.L. performed or contributed to the experiments on primary cell culture, with help from X.Z., H.L., and F.L. C.L., X.M. and C.D. performed the animal experiments. X.F., M.J., P.Y., and X.L. designed clinical protocols, reviewed clinical histories, selected and recruited study participants, and coordinated patient care teams to acquire profiled tissues. X.F. conceptualized and coordinated the study. P.W. and X.X wrote the manuscript. X.F., M.J., L.B., and W.Z. revised the manuscript. All authors reviewed and approved the manuscript.

## Data availability

All scRNA-seq and bulk RNA-seq data generated in this study have been deposited in Mendeley Data and will be made accessible upon acceptance.

## Code availability

All the codes related to the analysis are publicly available upon acceptance.

## Extended Data Figure Legends

**Extended Data Fig. 1:**
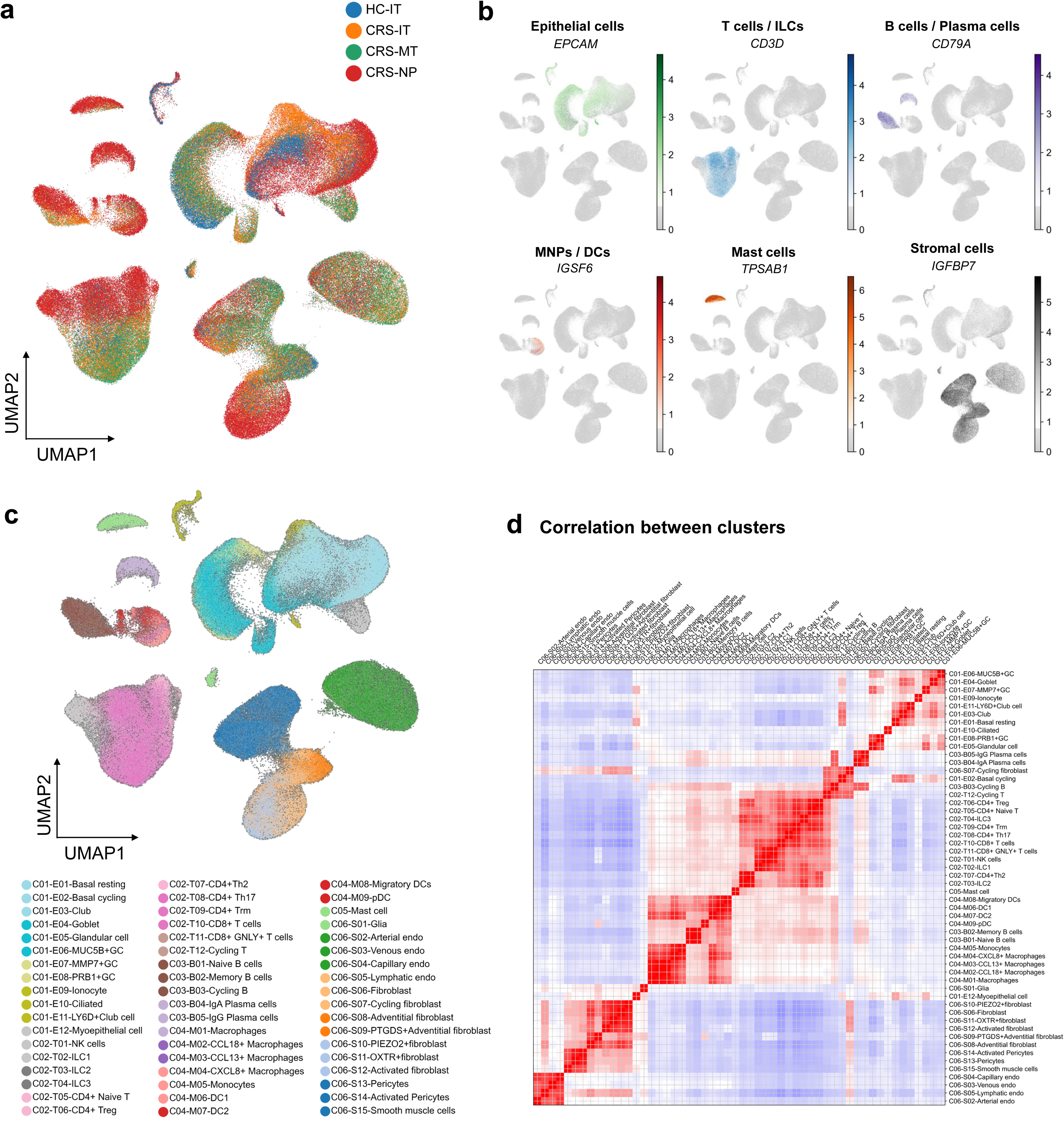
Landscape of the scRNA-seq data of CRSwNP. **a**, UMAP displaying total cells in the indicated anatomical regions. **b**, UMAP displaying expression of marker genes in the six cell compartments defined in Fig.1(**a**). **c**, UMAP displaying 54 cell subsets. **d**, Correlation analysis of the gene expression similarity of total cell subsets.

**Extended Data Fig. 2:**
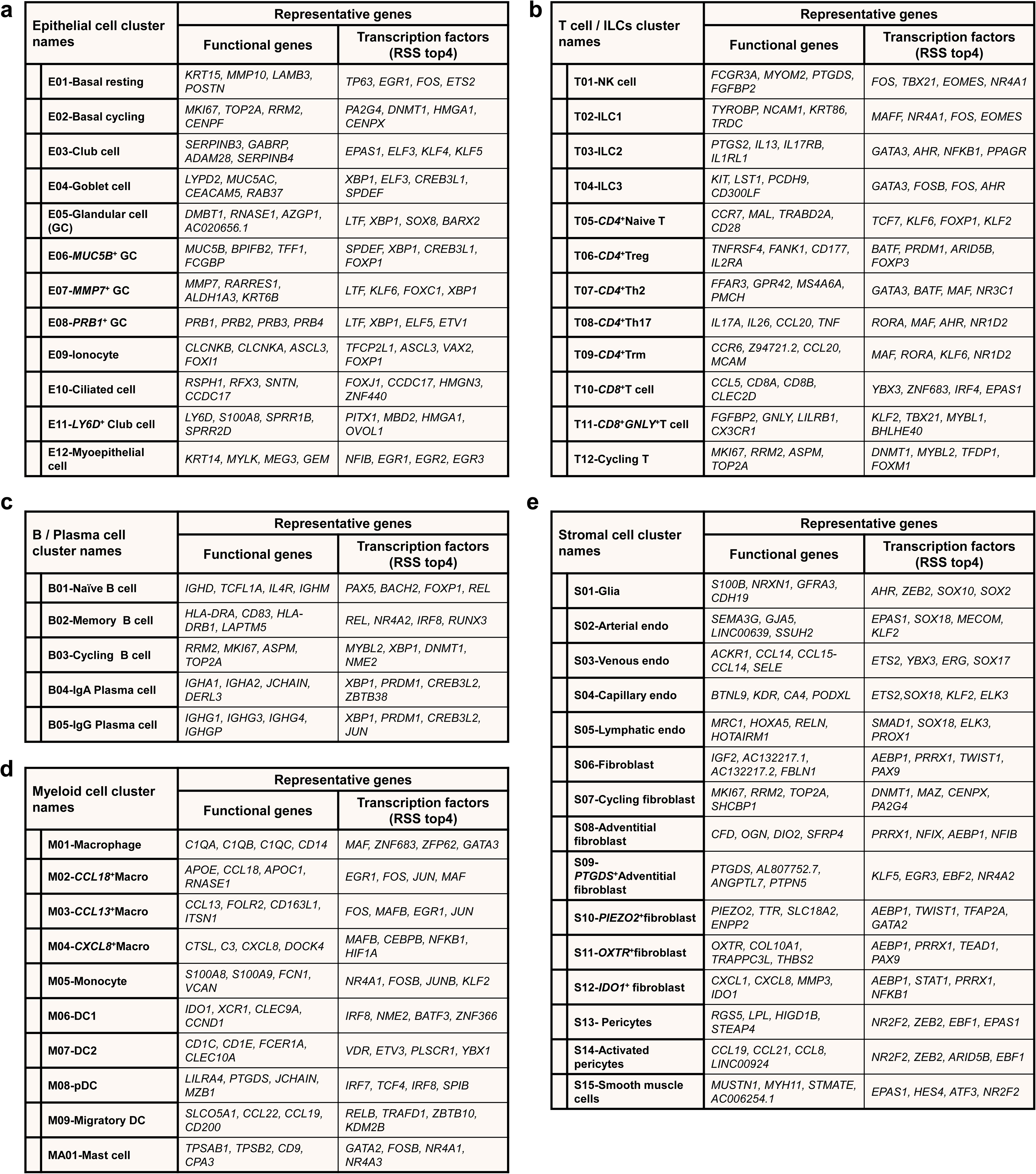
Summary of total cell subsets in nasal mucosa from patients with CRSwNP patients and healthy controls. (**a**-**d**) Overview of functional genes and key transcription factors of each cell subset. The four top genes are listed.

**Extended Data Fig. 3:**
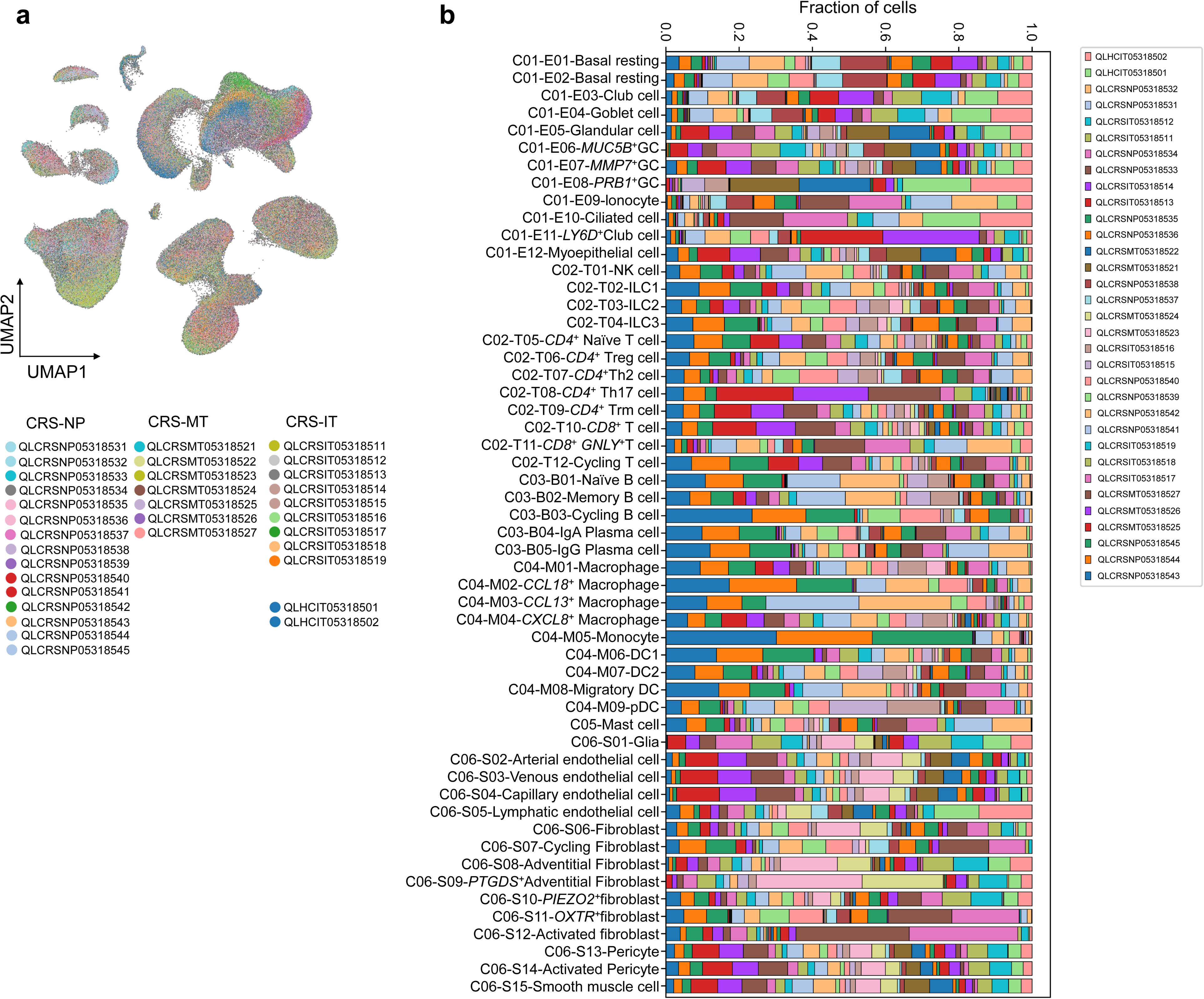
The distribution of samples in total cell subsets. **a**, UMAP embedding by patients with CRSwNP and healthy individuals. **b**, Bar plots showing the contributions of samples across total cell subsets.

**Extended Data Fig. 4:**
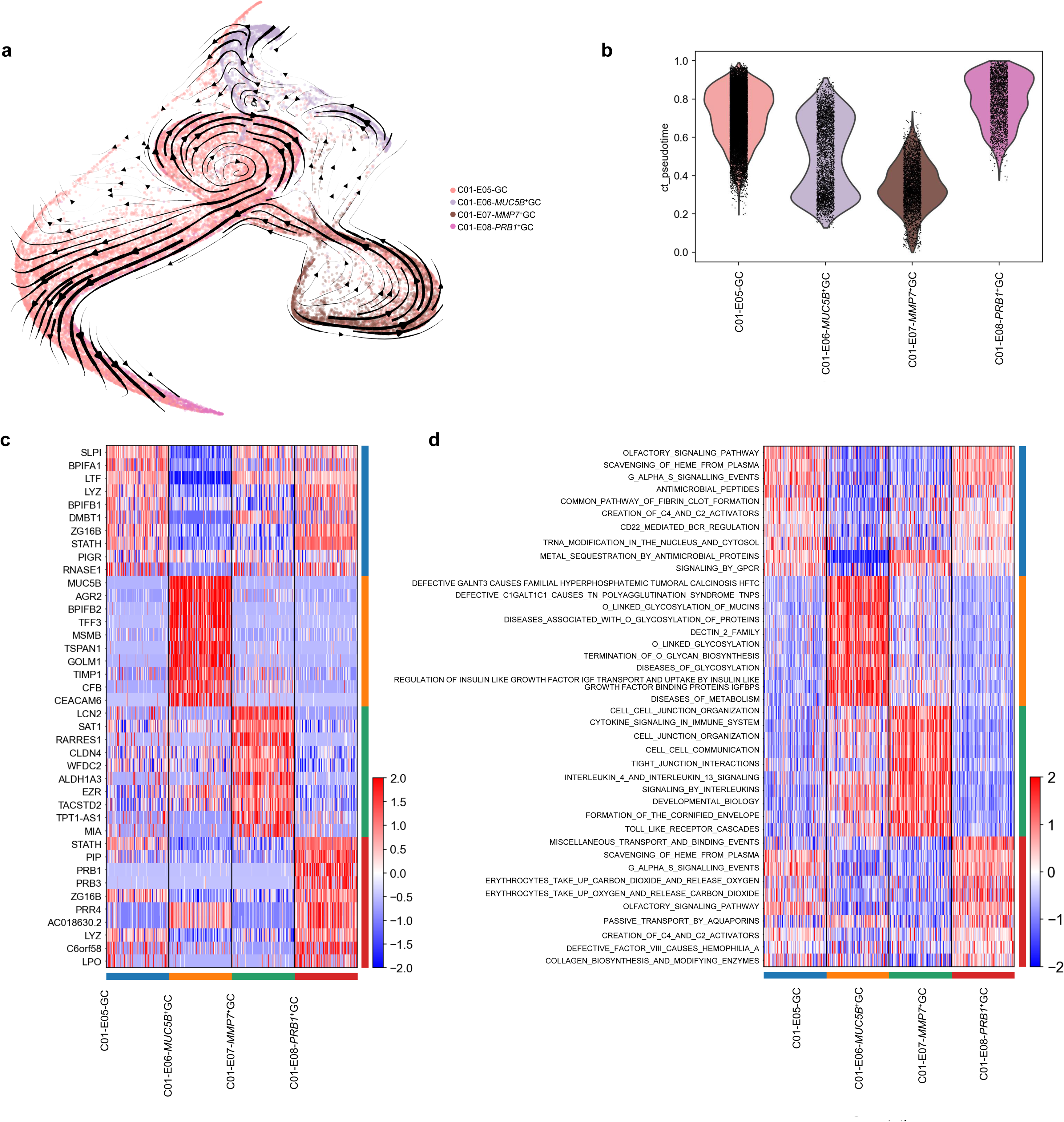
Analysis of glandular cells in nasal mucosa from CRSwNP patients and healthy controls. **a**, RNA velocity analysis based on RNA splicing information. **b**, Ct values of four glandular cell subsets calculated by pseudotime analysis. **c**, Heatmap displaying the expression of marker genes for four glandular cell subsets. **d**, Heatmap displaying enriched functional and signaling pathways for four glandular cell subsets.

**Extended Data Fig. 5:**
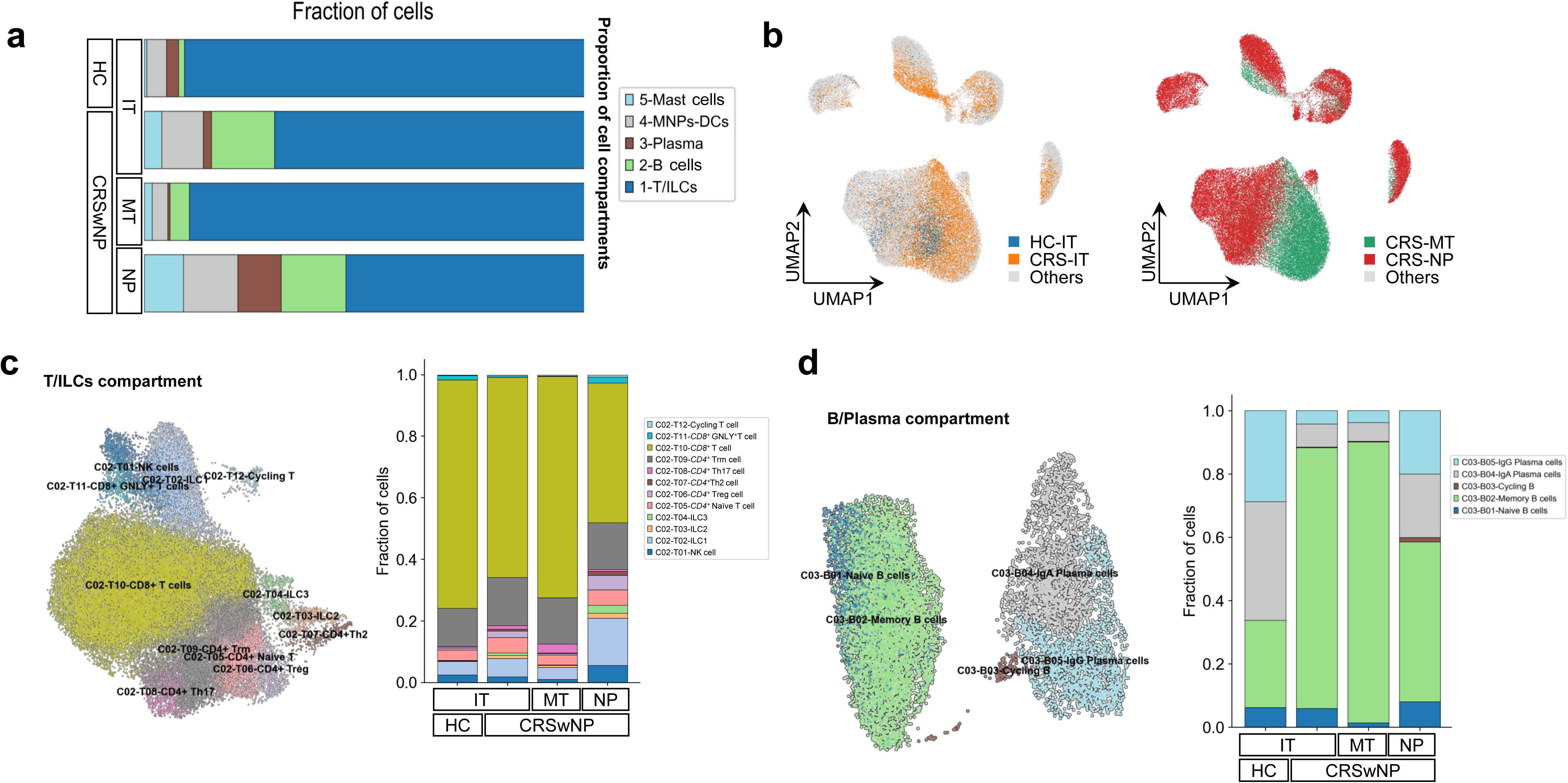
Sub-clustering of immune cells reiterates the inflammatory environment in neutrophilic CRSwNP. **a**, Bar plot displaying the proportions of five immune cell subsets from the indicated anatomical regions. **b**, UMAP displaying the distribution of immune cells in the indicated anatomical regions. **c**, UMAP displaying 12 T/ILCs subsets (left panel). The proportions of different T/ILCs cell subsets in the indicated anatomical regions (right panel). **d**, UMAP displaying 5 subsets of B cells and plasma cells (left panel). The proportions of cells in different B cell and plasma cell subsets in the indicated anatomical regions across different disease states (right panel).

**Extended Data Fig. 6:**
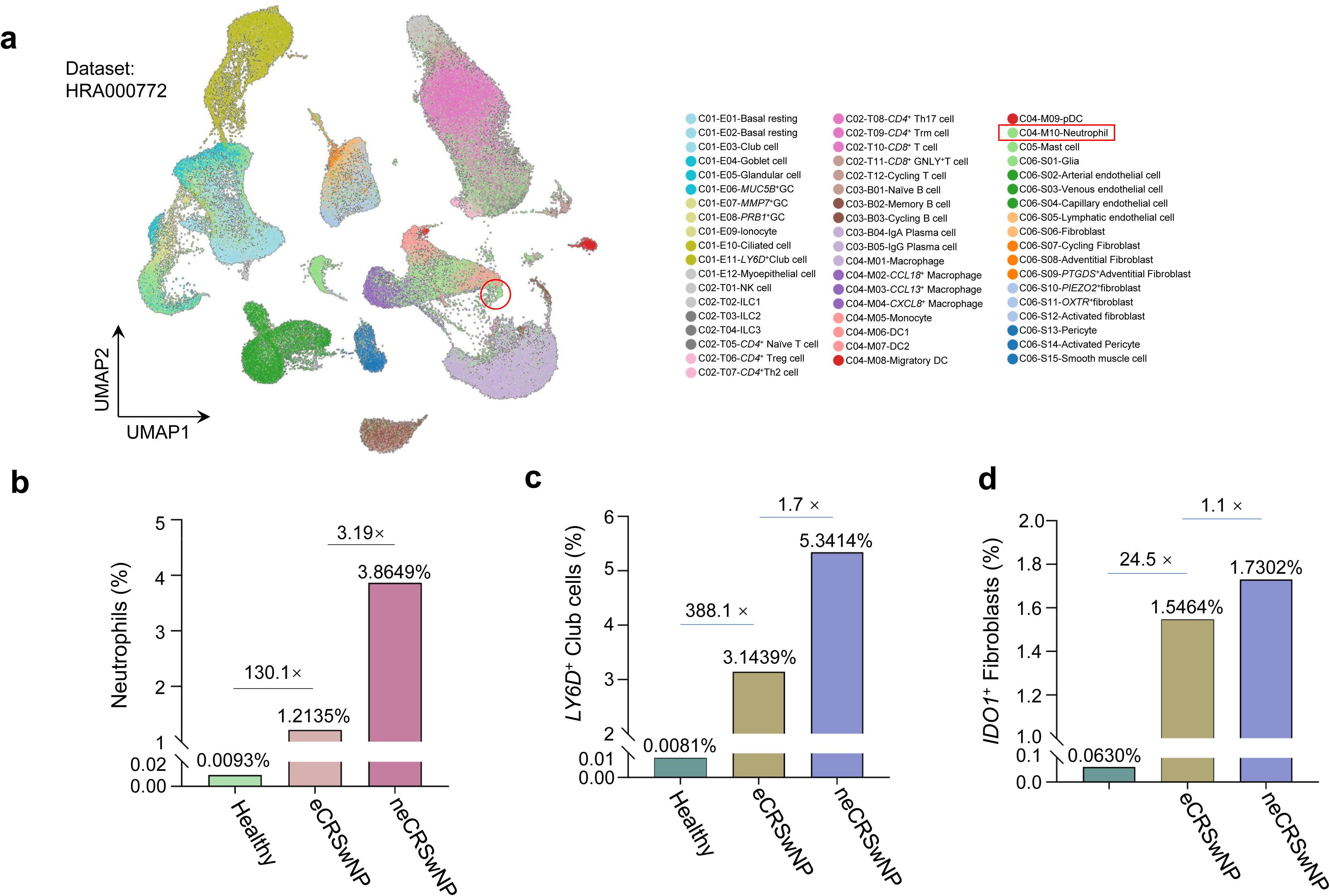
Analysis of the neutrophils, *LY6D*^+^ club cells, and *IDO1*^+^ fibroblasts in the scRNA-seq data of CRSwNP (HRA000772). **a**, UMAP displaying all cell subsets from the scRNA-seq dataset of CRSwNP (HRA000772). The enrichment of neutrophils is indicated by a red circle. **b**, Proportions of neutrophils in samples from eosinophilic, non-eosinophilic CRSwNP patients and healthy individuals. **c**, Proportions of *LY6D^+^* club cells in samples from eosinophilic, non-eosinophilic CRSwNP patients, and healthy individuals. **d**, Proportions of *IDO1*^+^ fibroblasts in samples from eosinophilic, non-eosinophilic CRSwNP patients, and healthy individuals.

**Extended Data Fig. 7:**
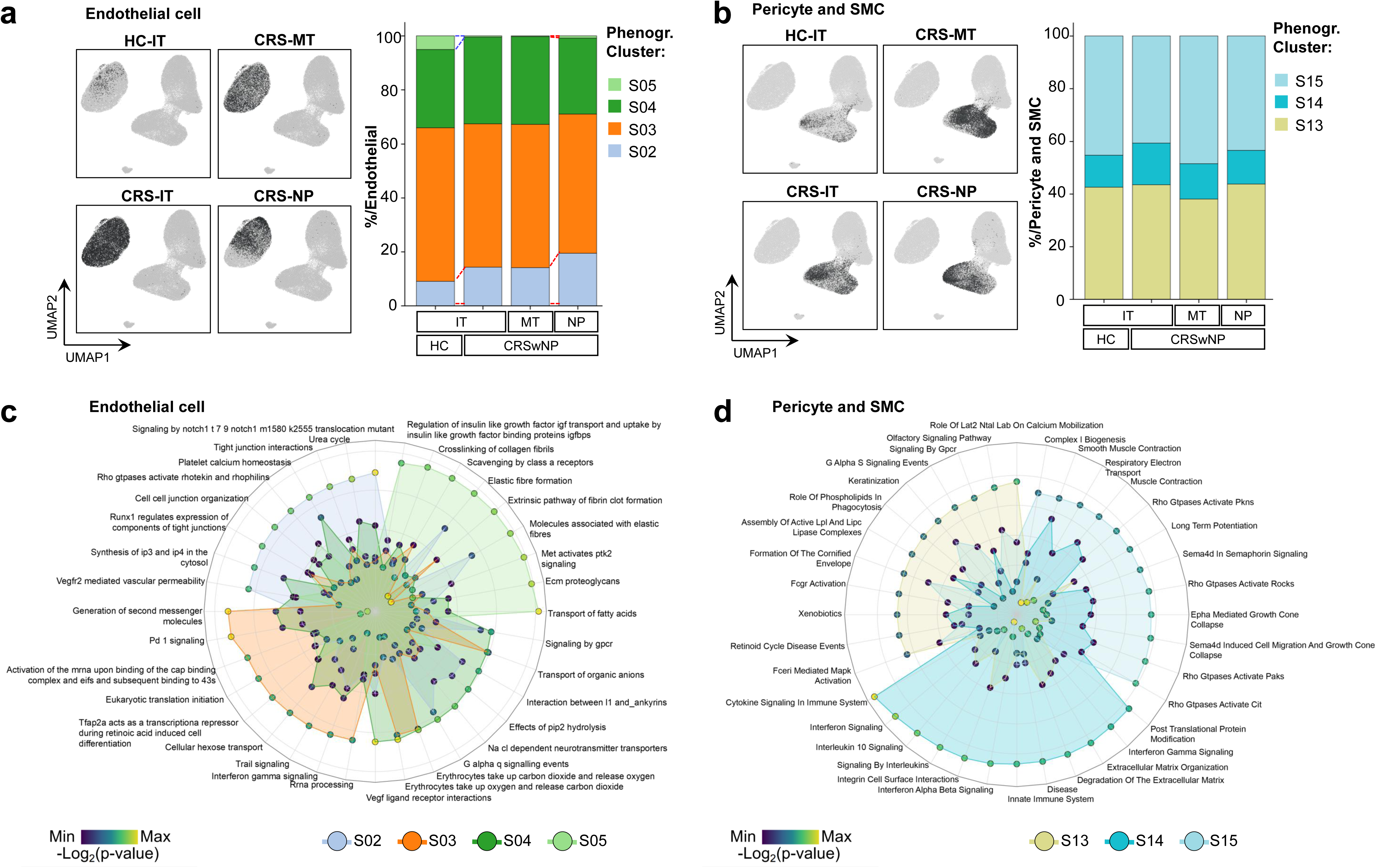
Endothelial cell, pericyte and smooth muscle cell subsets in nasal mucosa from CRSwNP patients and healthy controls. **a**, UMAP with stromal cell compartment displaying endothelial cells in the indicated anatomical regions (left panel). Bar plot depicting the proportions of the four endothelial cell subsets in the indicated anatomical regions (right panel). The numbering of the cell subsets is consistent with that in Fig. 1(g). **b**, UMAP with stromal cell compartment displaying pericytes and smooth muscle cells in the indicated anatomical regions (left panel). Bar plot depicting the distribution of three subsets of pericytes and smooth muscle cells in the indicated anatomical regions. The numbering of the cell subsets is consistent with that in Fig. 1(g). **c**, Pathway enrichment analysis displaying the enriched functional and signaling pathways for different endothelial cell subsets. **d**, Pathway enrichment analysis displaying the enriched functional and signaling pathways for different subsets of pericytes and smooth muscle cells.

**Extended Data Fig. 8:**
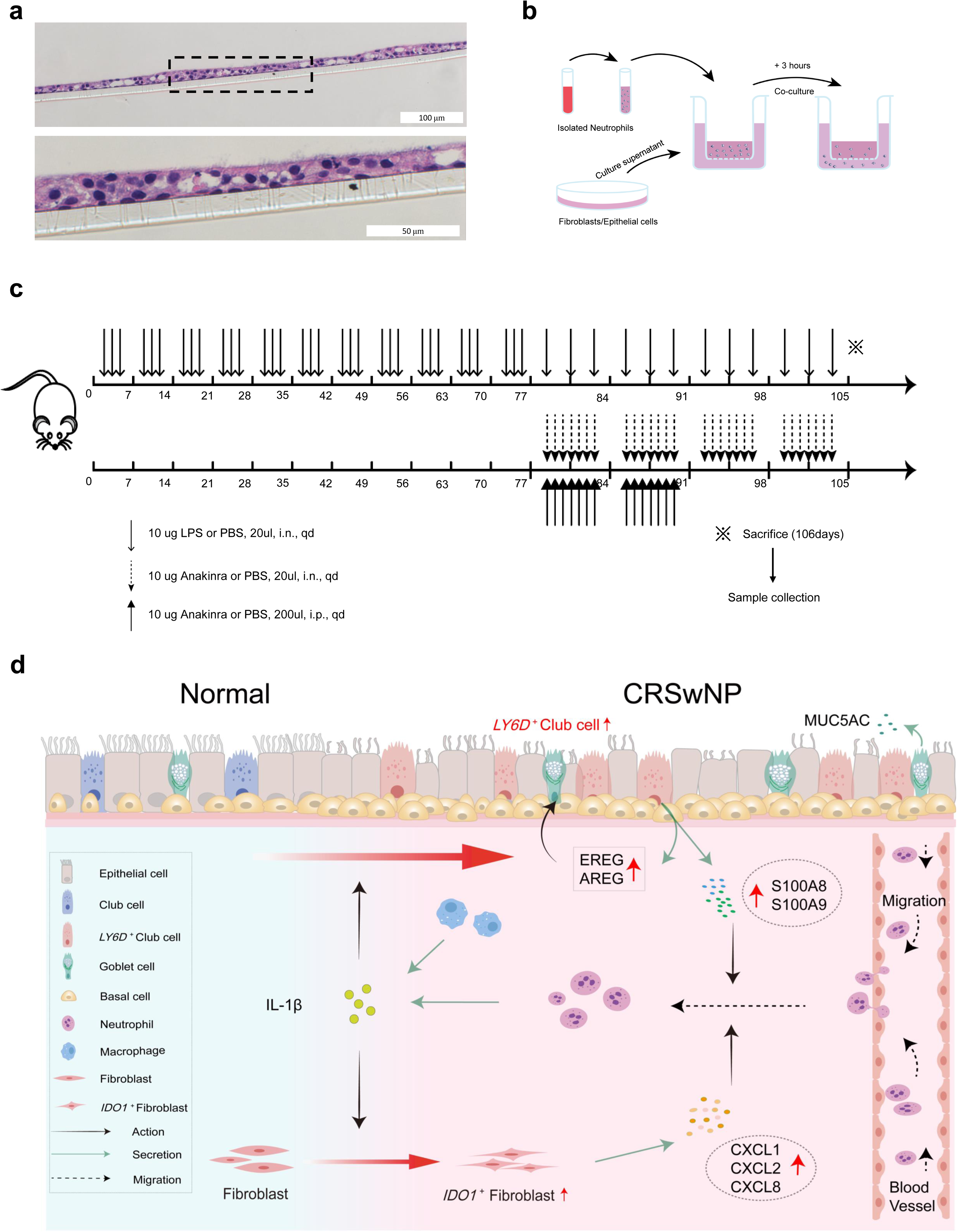
Graphical protocols and schematic diagram of the mechanism involved in this study. **a**, Representative H&E images of ALI-cultured HNEs showing the structure of differentiated epithelial cells, such as ciliated cells. Scale bar, 100 μm (upper), 50 μm (below). **B**, Schematic diagram of the neutrophil chemotaxis model. **c**, Graphical protocol for the establishment of a mouse model of neutrophilic chronic rhinosinusitis with nasal polyps induced by LPS. The protocol for anakinra administration was included. **d**, Graphical summary of new findings on CRSwNP pathways from this research. The graphical summary provides novel insights into the mechanism of neutrophil recruitment in the nasal mucosa of both CRSwNP patients and healthy individuals.

## Supplementary Materials

**Supplementary Table 1** Clinical characteristics of healthy control subjects and CRSwNP patients for this study

**Supplementary Table 2** Antibodies and other reagents in this study

## References

1. Gernez, Y., Tirouvanziam, R. & Chanez, P. Neutrophils in chronic inflammatory airway diseases: can we target them and how? Eur Respir J 35, 467–469 (2010).

2. Esther, C. R. et al. Metabolomic Evaluation of Neutrophilic Airway Inflammation in Cystic Fibrosis. Chest 148, 507–515 (2015).

3. Fokkens, W. J. et al. EPOS 2012: European position paper on rhinosinusitis and nasal polyps 2012. A summary for otorhinolaryngologists. Rhinology 50, 1–12 (2012).

4. Benjamin, M. R. et al. Clinical Characteristics of Patients with Chronic Rhinosinusitis without Nasal Polyps in an Academic Setting. J Allergy Clin Immunol Pract 7, 1010–1016 (2019).

5. Tomassen, P. et al. Inflammatory endotypes of chronic rhinosinusitis based on cluster analysis of biomarkers. J Allergy Clin Immunol 137, 1449–1456.e4 (2016).

6. Delemarre, T. & Bachert, C. Neutrophilic inflammation in chronic rhinosinusitis. Curr Opin Allergy Clin Immunol 23, 14–21 (2023).

7. Kobayashi, Y. The role of chemokines in neutrophil biology. Front Biosci 13, 2400–2407 (2008).

8. Wang, X. et al. Diversity of TH cytokine profiles in patients with chronic rhinosinusitis: A multicenter study in Europe, Asia, and Oceania. J Allergy Clin Immunol 138, 1344–1353 (2016).

9. Sumsion, J. S., Pulsipher, A. & Alt, J. A. Differential expression and role of S100 proteins in chronic rhinosinusitis. Curr Opin Allergy Clin Immunol 20, 14–22 (2020).

10. Wang, H. et al. The activation and function of IL-36γ in neutrophilic inflammation in chronic rhinosinusitis. J Allergy Clin Immunol 141, 1646–1658 (2018).

11. Palacios-García, J. et al. Role of Fibroblasts in Chronic Inflammatory Signalling in Chronic Rhinosinusitis with Nasal Polyps—A Systematic Review. J Clin Med 12, 3280 (2023).

12. Poposki, J. A. et al. Increased expression of the chemokine CCL23 in eosinophilic chronic rhinosinusitis with nasal polyps. J Allergy Clin Immunol 128, 73–81.e4 (2011).

13. Peng, Y. et al. Whole-transcriptome sequencing reveals heightened inflammation and defective host defence responses in chronic rhinosinusitis with nasal polyps. Eur Respir J 54, 1900732 (2019).

14. Rinia, A. B., Kostamo, K., Ebbens, F. A., van Drunen, C. M. & Fokkens, W. J. Nasal polyposis: a cellular-based approach to answering questions. Allergy 62, 348–358 (2007).

15. Ordovas-Montanes, J. et al. Allergic inflammatory memory in human respiratory epithelial progenitor cells. Nature 560, 649–654 (2018).

16. Morita, H., Moro, K. & Koyasu, S. Innate lymphoid cells in allergic and nonallergic inflammation. J Allergy Clin Immunol 138, 1253–1264 (2016).

17. Ma, J., et al. Single-cell analysis pinpoints distinct populations of cytotoxic CD4+ T cells and an IL-10+CD109+ TH2 cell population in nasal polyps. Sci Immunol 6, eabg6356 (2021).

18. Deprez, M. et al. A Single-Cell Atlas of the Human Healthy Airways. Am J Respir Crit Care Med 202, 1636–1645 (2020).

19. Delemarre, T. et al. A substantial neutrophilic inflammation as regular part of severe type 2 chronic rhinosinusitis with nasal polyps. J Allergy Clin Immunol 147, 179–188.e2 (2021).

20. Wang, W. et al. Single-cell profiling identifies mechanisms of inflammatory heterogeneity in chronic rhinosinusitis. Nat Immunol 23, 1484–1494 (2022).

21. Capucetti, A., Albano, F. & Bonecchi, R. Multiple Roles for Chemokines in Neutrophil Biology. Front Immunol 11, 1259 (2020).

22. Bonecchi, R. et al. Up-regulation of CCR1 and CCR3 and induction of chemotaxis to CC chemokines by IFN-gamma in human neutrophils. J Immunol 162, 474–479 (1999).

23. Nakatani, A. et al. S100A8 enhances IL-1β production from nasal epithelial cells in eosinophilic chronic rhinosinusitis. Allergol Int 72, 143–150 (2023).

24. Boruk, M. et al. Elevated S100A9 expression in chronic rhinosinusitis coincides with elevated MMP production and proliferation in vitro. Sci Rep 10, 16350 (2020).

25. Van Crombruggen, K., Vogl, T., Pérez-Novo, C., Holtappels, G. & Bachert, C. Differential release and deposition of S100A8/A9 proteins in inflamed upper airway tissue. Eur Respir J 47, 264–274 (2016).

26. Ryckman, C., Vandal, K., Rouleau, P., Talbot, M. & Tessier, P. A. Proinflammatory activities of S100: proteins S100A8, S100A9, and S100A8/A9 induce neutrophil chemotaxis and adhesion. J Immunol 170, 3233–3242 (2003).

27. Yoshisue, H. & Hasegawa, K. Effect of MMP/ADAM inhibitors on goblet cell hyperplasia in cultured human bronchial epithelial cells. Biosci Biotechnol Biochem 68, 2024–2031 (2004).

28. Casalino-Matsuda, S. M., Monzón, M. E. & Forteza, R. M. Epidermal growth factor receptor activation by epidermal growth factor mediates oxidant-induced goblet cell metaplasia in human airway epithelium. Am J Respir Cell Mol Biol 34, 581–591 (2006).

29. Mackay, C. D. A., Jadli, A. S., Fedak, P. W. M. & Patel, V. B. Adventitial Fibroblasts in Aortic Aneurysm: Unraveling Pathogenic Contributions to Vascular Disease. Diagnostics (Basel) 12, 871 (2022).

30. Stenmark, K. R. et al. The adventitia: Essential role in pulmonary vascular remodeling. Compr Physiol 1, 141–161 (2011).

31. Matsushima, K., Yang, D. & Oppenheim, J. J. Interleukin-8: An evolving chemokine. Cytokine 153, 155828 (2022).

32. He, Z. et al. Interleukin 1 beta and Matrix Metallopeptidase 3 Contribute to Development of Epidermal Growth Factor Receptor-Dependent Serrated Polyps in Mouse Cecum. Gastroenterology 157, 1572–1583.e8 (2019).

33. Knight, D. A. et al. Leukemia inhibitory factor (LIF) and LIF receptor in human lung. Distribution and regulation of LIF release. Am J Respir Cell Mol Biol 20, 834–841 (1999).

34. Dinarello, C. A. Overview of the IL-1 family in innate inflammation and acquired immunity. Immunol Rev 281, 8–27 (2018).

35. Anti-Interleukin-1 Beta/Tumor Necrosis Factor-Alpha IgY Antibodies Reduce Pathological Allergic Responses in Guinea Pigs with Allergic Rhinitis - PubMed. https://pubmed.ncbi.nlm.nih.gov/27046957/.

36. Wang, S. et al. Establishment of a mouse model of lipopolysaccharide-induced neutrophilic nasal polyps. Exp Ther Med 14, 5275–5282 (2017).

37. Jang, Y. et al. Anakinra treatment for refractory cerebral autoinflammatory responses. Ann Clin Transl Neurol 9, 91–97 (2022).

38. Cavalli, G. & Dinarello, C. A. Anakinra Therapy for Non-cancer Inflammatory Diseases. Front Pharmacol 9, 1157 (2018).

39. Eldridge, M. W. & Peden, D. B. Allergen provocation augments endotoxin-induced nasal inflammation in subjects with atopic asthma. J Allergy Clin Immunol 105, 475–481 (2000).

40. Wei, Y. et al. Activated pyrin domain containing 3 (NLRP3) inflammasome in neutrophilic chronic rhinosinusitis with nasal polyps (CRSwNP). J Allergy Clin Immunol 145, 1002–1005.e16 (2020).

41. Ruan, J.-W. et al. Characterizing the Neutrophilic Inflammation in Chronic Rhinosinusitis With Nasal Polyps. Front Cell Dev Biol 9, 793073 (2021).

42. Ding, G. Q., Zheng, C. Q. & Bagga, S. S. Up-regulation of the mucosal epidermal growth factor receptor gene in chronic rhinosinusitis and nasal polyposis. Arch Otolaryngol Head Neck Surg 133, 1097–1103 (2007).

43. de Boer, W. I. et al. Expression of epidermal growth factors and their receptors in the bronchial epithelium of subjects with chronic obstructive pulmonary disease. Am J Clin Pathol 125, 184–192 (2006).

44. Cheng, W.-L. et al. The Role of EREG/EGFR Pathway in Tumor Progression. Int J Mol Sci 22, 12828 (2021).

45. Schiwon, M. et al. Crosstalk between sentinel and helper macrophages permits neutrophil migration into infected uroepithelium. Cell 156, 456–468 (2014).

46. Kim, N. D. & Luster, A. D. The role of tissue resident cells in neutrophil recruitment. Trends Immunol 36, 547–555 (2015).

47. Kikuchi, I. et al. Eosinophil trans-basement membrane migration induced by interleukin-8 and neutrophils. Am J Respir Cell Mol Biol 34, 760–765 (2006).

48. Moore, W. C. et al. Sputum neutrophil counts are associated with more severe asthma phenotypes using cluster analysis. J Allergy Clin Immunol 133, 1557–1563.e5 (2014).

49. Tai, J., Han, M. & Kim, T. H. Therapeutic Strategies of Biologics in Chronic Rhinosinusitis: Current Options and Future Targets. Int J Mol Sci 23, 5523 (2022).

50. Abadie, B. Q. & Cremer, P. C. Interleukin-1 Antagonists for the Treatment of Recurrent Pericarditis. BioDrugs 36, 459–472 (2022).

51. Fokkens, W. J. et al. European Position Paper on Rhinosinusitis and Nasal Polyps 2020. Rhinology 58, 1–464 (2020).

52. Wolf, F. A., Angerer, P. & Theis, F. J. SCANPY: large-scale single-cell gene expression data analysis. Genome Biol 19, 15 (2018).

53. Wolock, S. L., Lopez, R. & Klein, A. M. Scrublet: Computational Identification of Cell Doublets in Single-Cell Transcriptomic Data. Cell Syst 8, 281–291.e9 (2019).

54. Tirosh, I. et al. Dissecting the multicellular ecosystem of metastatic melanoma by single-cell RNA-seq. Science 352, 189–196 (2016).

55. Polański, K. et al. BBKNN: fast batch alignment of single cell transcriptomes. Bioinformatics 36, 964–965 (2020).

56. Traag, V. A., Waltman, L. & van Eck, N. J. From Louvain to Leiden: guaranteeing well-connected communities. Sci Rep 9, 5233 (2019).

57. Becht, E. et al. Dimensionality reduction for visualizing single-cell data using UMAP. Nat Biotechnol (2018) doi:10.1038/nbt.4314.

58. Van de Sande, B. et al. A scalable SCENIC workflow for single-cell gene regulatory network analysis. Nat Protoc 15, 2247–2276 (2020).

59. Aibar, S. et al. SCENIC: single-cell regulatory network inference and clustering. Nat Methods 14, 1083–1086 (2017).

60. Lambert, S. A. et al. The Human Transcription Factors. Cell 172, 650–665 (2018).

61. He, D. et al. Alevin-fry unlocks rapid, accurate and memory-frugal quantification of single-cell RNA-seq data. Nat Methods 19, 316–322 (2022).

62. Domínguez Conde, C., et al. Cross-tissue immune cell analysis reveals tissue-specific features in humans. Science 376, eabl5197 (2022).

63. La Manno, G. et al. RNA velocity of single cells. Nature 560, 494–498 (2018).

64. Lange, M. et al. CellRank for directed single-cell fate mapping. Nat Methods 19, 159–170 (2022).

65. Croft, D. et al. Reactome: a database of reactions, pathways and biological processes. Nucleic Acids Res 39, D691–697 (2011).

66. Orchard, S. et al. Protein interaction data curation: the International Molecular Exchange (IMEx) consortium. Nat Methods 9, 345–350 (2012).

67. Orchard, S. et al. The MIntAct project--IntAct as a common curation platform for 11 molecular interaction databases. Nucleic Acids Res 42, D358–363 (2014).

68. Breuer, K. et al. InnateDB: systems biology of innate immunity and beyond--recent updates and continuing curation. Nucleic Acids Res 41, D1228–1233 (2013).

69. Licata, L. et al. MINT, the molecular interaction database: 2012 update. Nucleic Acids Res 40, D857–861 (2012).

70. Brown, K. R. & Jurisica, I. Unequal evolutionary conservation of human protein interactions in interologous networks. Genome Biol 8, R95 (2007).

71. Aran, D., Hu, Z. & Butte, A. J. xCell: digitally portraying the tissue cellular heterogeneity landscape. Genome Biol 18, 220 (2017).

72. Beppu, A. K. et al. Epithelial plasticity and innate immune activation promote lung tissue remodeling following respiratory viral infection. Nat Commun 14, 5814 (2023).

73. Chen, W. et al. Over-expression of CRTH2 indicates eosinophilic inflammation and poor prognosis in recurrent nasal polyps. Front Immunol 13, 1046426 (2022).

