## Supplemental Tables for "Transdifferentiation of epithelial cells and fibroblasts induced by IL-1β fuels neutrophil recruitment in chronic rhinosinusitis": Supplementary Table1.pdf

**Supplementary Table 1: Clinical characteristics of healthy control subjects and CRSwNP patients in this study**

| | Healthy Controls | CRSwNP | *P* value |
|---|---|---|---|
| | n or mean±SD | n or mean±SD | |
| **Number of Subjects** | 38 | 47 | |
| **Gender（Female/Male）** | 9/29 | 14/33 | 0.53 |
| **Age** | 39.13±13.14 | 44.30±12.40 | 0.07 |
| **With Allergic Rhinitis** | 0 | 10 | <0.01 |
| **With Asthma** | 0 | 8 | <0.01 |
| **Smoking** | 14 | 18 | 0.89 |
| **White blood cell (WBC)** | 6.29±1.38 | 6.71±1.43 | 0.20 |
| **Blood Neutrophil Percentage (%)** | 53.02±7.98 | 52.6±7.99 | 0.92 |
| **Blood Lymphocyte Percentage (%)** | 36.73±7.72 | 35.01±8.77 | 0.26 |
| **Blood Eosinophil Percentage (%)** | 2.34±1.70 | 5.34±3.60 | <0.01 |
| **Blood Basophil Percentage (%)** | 0.54±0.21 | 0.74±0.77 | 0.12 |
| **Blood Monocyte Percentage (%)** | 7.36±1.57 | 7.09±1.74 | 0.48 |
| **Blood Neutrophil Count (10^9/L)** | 3.39±1.23 | 3.56±1.05 | 0.48 |
| **Blood Lymphocyte Count (10^9/L)** | 2.26±0.49 | 2.28±0.65 | 0.94 |
| **Blood Eosinophil Count (10^9/L)** | 0.13±0.10 | 0.35±0.22 | <0.01 |
| **Blood Basophil Count (10^9/L)** | 0.04±0.02 | 0.05±0.04 | 0.28 |
| **Blood Monocyte Count (10^9/L)** | 0.46±0.15 | 0.48±0.11 | 0.64 |
