## Supplemental Tables for "Transdifferentiation of epithelial cells and fibroblasts induced by IL-1β fuels neutrophil recruitment in chronic rhinosinusitis": Supplementary Table 2(1).pdf

**Supplementary Table 1. Antibodies and other reagents in this study.**

| Reagents | Company | Catalog Number | Dilution |
|---|---|---|---|
| Anti-LY6D | Sigma | HPA024755 | 1/2000 |
| Anti-LY6D | Proteintech | 17361-1-AP | 1/100 |
| Anti-SPRR1B | Proteintech | 11959-1-AP | 1/100 |
| Anti-S100A8 | Thermo Fisher | MA5-17621 | 1/100 |
| Anti-KRT13 | Abcam | ab16112 | 1/1000 |
| Anti-KRT13 | Proteintech | 10164-2-AP | 1/200 |
| Anti-CXCL8 | Santa Cruz | sc-376750 | 1/200 |
| Anti-IDO1 | Abcam | ab211071 | 1/1000 |
| Anti-IDO1 | Santa Cruz | sc-53978 | 1/200 |
| Anti-COL1A2 | Santa Cruz | sc-393573 | 1/200 |
| Anti-MPO | Abcam | ab208670 | 1/1000 |
| Anti-IL-1β | Bioss | bs-0812R | 1/200 |
| CoraLite594-conjugated Goat Anti-Mouse | Proteintech | SA00013-3 | 1/500 |
| CoraLite488-conjugated Goat Anti-Rabbit | Proteintech | SA00013-2 | 1/500 |
| AR6 Buffer | Akoya | FP1498 | |
| Opal 520 Reagent | Akoya | FP1487A | |
| Opal 570 Reagent | Akoya | FP1488A | |
| Opal 650 Reagent | Akoya | FP1496A | |
| Spectral 4',6-diamidino-2-phenylindole（DAPI） | Akoya | FP1490A | |
| Antibody Diluent | Akoya | ARD1001EA | |
| PolyHRP Broad Spectrum | Akoya | ARH1001EA | |
| 1X Plus Amplification Diluent | Akoya | AR600125ML | |
| Diaminobenzidine Horseradish Peroxidase Color Development Kit | ZSGB-Bio | PV-9000 | |
| Human IL-8 ELISA Kit | 4A Biotech | CHE0011 | |
| Mouse IL-8 ELISA Kit | 4A Biotech | CME0008 | |
| Mouse TNFα ELISA Kit | 4A Biotech | CME0004 | |
| Human S100A8/S100A9 Heterodimer Quantikine ELISA Kit | R&D SYSTEM | DS8900 | |
